# Development and Validation of Subject-Specific 3D Human Head Models Based on a Nonlinear Visco-Hyperelastic Constitutive Framework

**DOI:** 10.1101/2021.10.28.466370

**Authors:** Kshitiz Upadhyay, Ahmed Alshareef, Andrew K. Knutsen, Curtis L. Johnson, Aaron Carass, Philip V. Bayly, K.T. Ramesh

**Affiliations:** Hopkins Extreme Materials Institute, Johns Hopkins University, Baltimore, MD 21218, USA; Department of Mechanical Engineering, Johns Hopkins University, Baltimore, MD 21218, USA; Department of Electrical and Computer Engineering, Johns Hopkins University, Baltimore, MD 21218, USA; Center for Neuroscience and Regenerative Medicine, The Henry M. Jackson Foundation for the Advancement of Military Medicine, Bethesda, MD 20814, USA; Department of Biomedical Engineering, University of Delaware, Newark, DE 19716, USA; Mechanical Engineering and Materials Science, Washington University in St. Louis, St. Louis, MO 63130, USA

**Keywords:** Subject-specific simulations, Traumatic Brain Injury (TBI), Visco-hyperelasticity, Viscous dissipation potential, Head injury model, Magnetic resonance elastography

## Abstract

Computational models of the human head are promising tools for the study and prediction of traumatic brain injuries (TBIs). Most available head models are developed using inputs (i.e., head geometry, material properties, and boundary conditions) derived from ex-vivo experiments on cadavers or animals and employ linear viscoelasticity (LVE)-based constitutive models, which leads to high uncertainty and poor accuracy in capturing the nonlinear response of brain tissue under impulsive loading conditions. To resolve these issues, a framework for the development of fully subject-specific 3D human head models is proposed, in which model inputs are derived from the same living human subject using a comprehensive in-vivo brain imaging protocol, and the viscous dissipation-based visco-hyperelastic constitutive modeling framework is employed. Specifically, brain tissue material properties are derived from in-vivo magnetic resonance elastography (MRE), and full-field strain-response of brain under rapid rotational acceleration is obtained from tagged MRI, which is used for model validation. The constitutive model comprises the Ogden hyperelastic strain energy density and the Upadhyay-Subhash-Spearot viscous dissipation potential. The simulated strain-response is compared with experimental data and with predictions from subject-specific models employing two commonly used LVE-based constitutive models, using a rigorous validation procedure that evaluates agreement in spatial strain distribution, temporal strain evolution, and differences in maximum values of peak and average strain. Results show that the head model developed in this work reasonably captures 3D brain dynamics, and when compared to LVE-based models, provides improvements in the prediction of peak strains and temporal strain evolution.

## 1. Introduction

Traumatic brain injury (TBI) is one of the leading causes of death and disability in the United States, with the latest data showing nearly 61,000 TBI-related fatalities in 2019^[1]^. TBI is often caused by external forces on and rapid motions of the head. Computational human head models, which can simulate the response of the brain under rapid loading conditions, play an important role in the understanding and prediction of TBI^[2]^.

A number of computational head models have been developed over the past two decades^[3]^, each comprising three basic inputs: model geometry, brain tissue material properties, and boundary or loading conditions. Typically, model geometry either represents an “average” head^[4]^ or is obtained from imaging data of a representative subject^[5]^. Material properties are generally derived from ex-vivo or in-vitro experiments on brain tissues of animals or cadavers^[6]^. Finally, boundary or loading conditions from an experiment^[7,8]^ are applied to the head model. The resulting simulated response is then compared with the available experimental data for model validation. As brain morphometry can vary significantly both within and across different animal species, and tissue material properties can change between in-vivo and ex-vivo conditions^[9–11]^, these models are associated with significant uncertainty in the simulated response (and consequently, in the reported validation scores).

Existing head models often suffer from poor accuracy in comparison to experimental response^[4,5,12,13]^. A major factor that determines computational model accuracy is the underlying constitutive model used to define brain tissue behavior. Brain tissues exhibit features such as^[11,14,15]^: large deformations, nonlinear stress versus strain response, strain stiffening, strain rate sensitivity, and a rapidly increasing shear modulus with superimposed compression but not with superimposed tension, among others. These behaviors are particularly important when the head undergoes a rapid motion that leads to large strains and high strain rates, and thus constitutive models employed in head injury models should be able to capture these features to ensure reasonable accuracy. However, a majority of the available head models employ a linear viscoelastic (LVE) constitutive framework^[13,16,17]^, which assumes a linear elastic stress-strain response and a linear relaxation response. Despite its popularity, the LVE framework is not capable of capturing most of the characteristic features of brain tissue under large deformations and at high strain rates^[11,14,18]^, and thus is limited in capturing the response of the brain to injurious impact.

In addition to LVE models, two hereditary integral-based visco-hyperelastic constitutive models^[19]^ have also been used in computational head models: (i) quasi-linear viscoelastic (QLV)^[6,20]^ models, and (ii) models that assume an additive split of total stress into nonlinear hyperelasticity and linear viscoelasticity^[5,21]^. While these models can capture nonlinearity in the elastic part of the stress-strain response and are thus improvements, they assume a linear viscous part (or relaxation behavior). For human brain tissue, a number of previous studies have experimentally established that such a linear relaxation assumption only holds in the regime of up to approximately 1% strain^[22–24]^. Thus, applying these models to highly nonlinear brain tissue under rapid impulsive loading conditions can still lead to errors in the model predictions. A fully nonlinear constitutive framework is needed to capture the complexities of the mechanical response of brain in head models intended for use in TBI research.

In this study, fully subject-specific computational 3D human head models are developed, in which all the three basic inputs of model geometry, material properties, and boundary condition are derived from the same living human subject. This approach minimizes uncertainty due to the differences in brain structure and properties among the human population. Advanced in-vivo biomedical imaging techniques are employed for model development: magnetic resonance imaging (MRI), susceptibility weighted imaging (SWI), diffusion tensor imaging (DTI), magnetic resonance elastography (MRE), and tagged MRI (tMRI). A fully nonlinear viscous dissipation-based visco-hyperelastic constitutive model^[19,25]^ based on the Ogden hyperelastic strain energy density^[26]^ and the Upadhyay-Subhash-Spearot (USS) viscous dissipation potential^[19]^ is employed to capture the complex, nonlinear mechanical response of brain tissues. Constitutive model parameters are derived from the storage and loss moduli provided by MRE and then extended using a new hybrid model parametrization procedure. The subject-specific 3D head model is used to simulate brain deformations under mild rotational acceleration to the head, and the resulting computed strain fields are compared with experimentally observed strain fields from tMRI under the same loading condition. Finally, the predicted strain-response is compared with that from the corresponding head models that use the LVE and the Ogden model-based linear visco-hyperelastic constitutive models^[5,27]^.

## 2. Experimental Methods

A comprehensive in-vivo brain imaging protocol is conducted in this study on three healthy adult male subjects (age: 28, 31, and 46 years-old) to obtain inputs for the development of the computational model, as illustrated in Figure 1. Complete dsetails on the data acquisition, subject recruitment, and study approval are provided elsewhere^[28,29]^. Briefly, volumetric anatomical brain images from T_1_- and T_2_-weighted MRI and SWI are acquired to obtain the 3D model geometry (Fig. 1(a)). MRI images (spatial resolution: 0.8 mm isotropic) are skull-stripped and segmented into different anatomical substructures using a multi-atlas segmentation and cortical reconstruction algorithm^[30]^. The segmented structures are: deep gray matter, cortical gray matter, cerebral white matter, cerebellar gray matter, cerebellar white matter, brainstem, ventricles, and cerebrospinal fluid (CSF). From these images and segmentation labels, subarachnoid space (SAS) is automatically defined using a method described by Glaister et al.^[31]^. Further, falx and tentorium are reconstructed from T1-weighted and SWI images using a fast-marching multi-atlas-based segmentation method^[32]^. Corpus callosum is delineated from cerebral white matter using a semi-automatic 3D segmentation software, ITK-SNAP^[33]^, as described by Shiino et al.^[34]^. The fully segmented brain image is then rigidly registered to the brain biomechanics data (tMRI/MRE space) using the ANTs software package^[35]^. ANTs is also used to reduce the spatial image resolution to 1.5 x 1.5 x 1.5 mm^3^ to match the resolution of the tMRI data. Finally, a dilatation of the brain mask obtained via skull stripping with MONSTR^[36]^ is used to generate an artificial skull. The different substructures of the computational head model are illustrated in Fig. 1(b).

**Figure 1.**
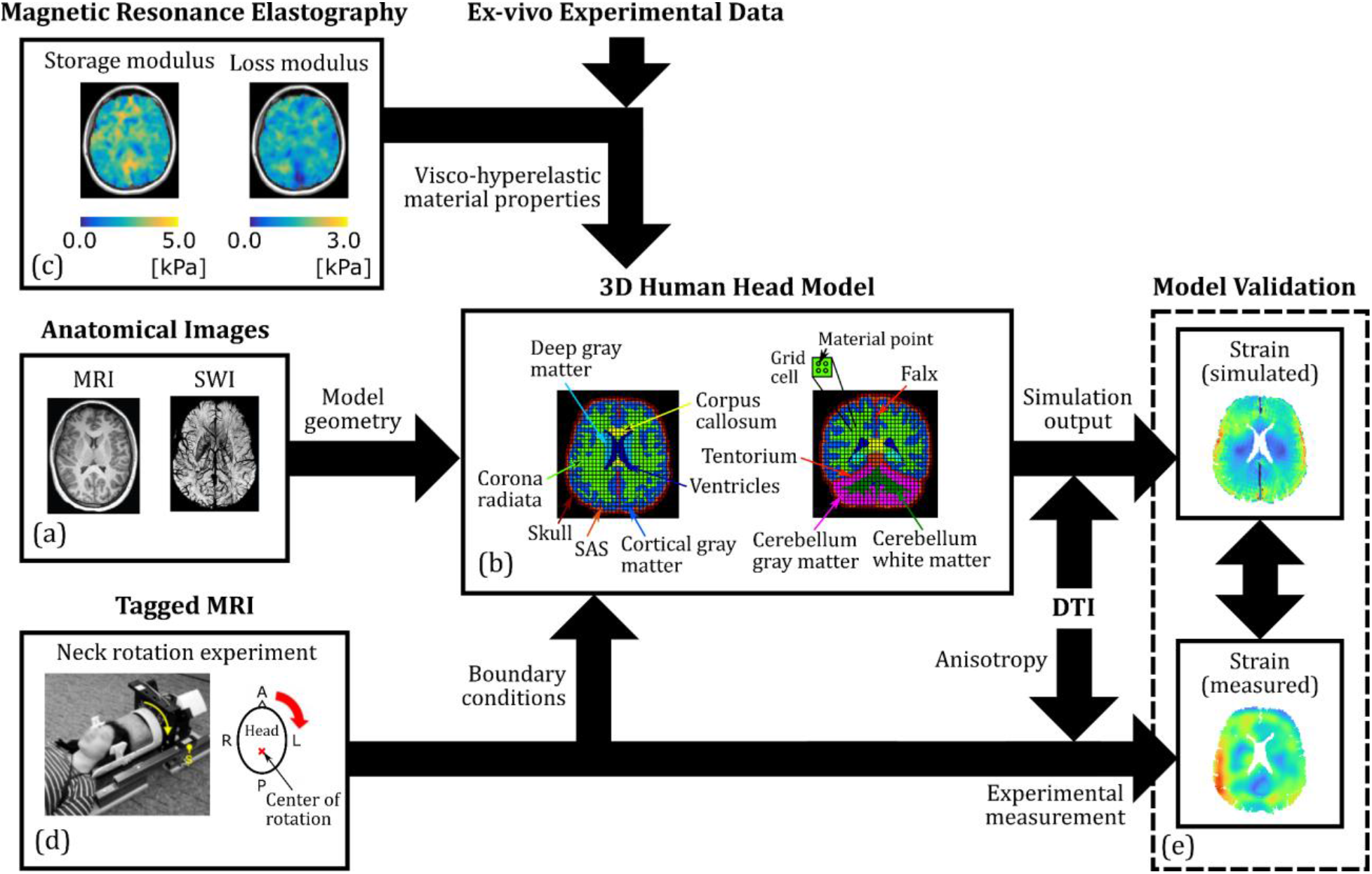
Flowchart of the development of subject-specific computational human head models: (a) Segmented and processed anatomical images from MRI (T1- and T2-weighted) and SWI provide the 3D geometry for (b) the head model; (c) MRE provides frequency-dependent storage and loss moduli, which are combined with ex-vivo stress-strain data from the literature to calibrate visco-hyperelastic material properties of major brain substructures. The MPM-based head model is used to simulate a (d) neck rotation experiment. Tagged MRI of this experiment yields full-field strain data for (e) computational model validation.

MRE^[37]^ is performed to derive in-vivo material properties of the different brain substructures (Fig. 1(c)). From imaged brain tissue displacement data, an advanced nonlinear inversion algorithm (NLI)^[38,39]^ is employed to estimate spatially-resolved full-field 3D maps of the storage modulus and loss modulus at 2.5 mm isotropic resolution. Like MRI images, MRE maps are rigidly registered to the tMRI data using ANTs, which also changes the resolution to 1.5 mm isotropic. Three separate scans are conducted at different actuation frequencies (i.e., 30, 50 and 70 Hz) to capture the frequency-dependent linear viscoelastic material response in the small-strain regime (strains generated by the MRE actuator system are on the order of ~10^−4^–10^−3^ mm/mm). This data, along with the ex-vivo large deformation stress versus strain response of brain tissues from the literature are utilized to calibrate the visco-hyperelastic material parameters for the head model as described in following sections.

Finally, multi-slice tMRI is employed in conjunction with the harmonic phase finite element method (HARP-FE)^[40]^ to acquire full-field 3D strains from neck rotation experiments, as illustrated in Figs. 1(d-e) and described by Knutsen et al.^[41]^. Briefly, in this experiment, a controlled non-injurious impulsive loading is applied on the subject’s head, which rotates in the axial plane about the inferior-superior (I/S) axis (center of rotation roughly passes through the brain stem). The loading input to the head is measured using an angular position sensor; figures 2(a) and 2(b) show the angular velocity and acceleration versus time plots, respectively, for one subject, S01 (31 years-old male). A typical range of peak angular head acceleration is approximately 150-350 rad/s^2^ and that of peak angular velocity is 1.5-3.5 rad/s. Loading input from this experiment is used as boundary condition to the computational head model, which outputs time-varying 3D Green-Lagrange (G-L) strain tensor fields. These simulated strain fields are compared with the experimentally measured strain fields to validate the head model, as illustrated in Fig. 1(e). In the HARP-FE post-processing phase, the raw tMRI data (spatial resolution: 2 mm^2^ axial in-plane, 8 mm out-of-plane) is resampled to a 1.5 mm isotropic grid, resulting in full-field strains appropriate for comparison with the simulated strain data. The temporal resolution of tMRI is 18 ms, and the total measurement duration is 189 ms.

**Figure 2.**
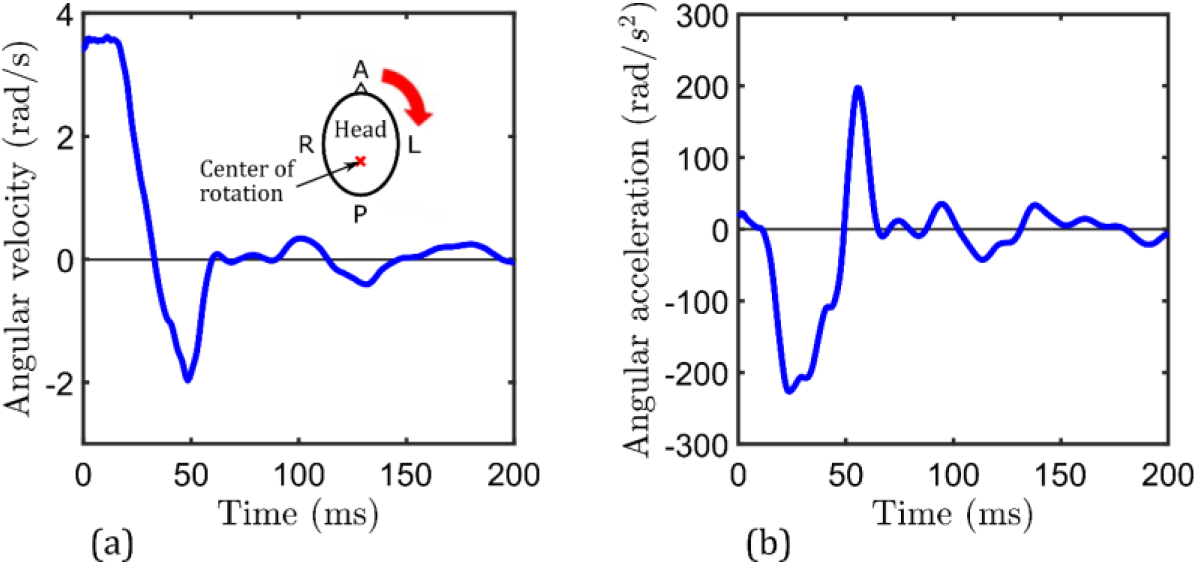
Head kinematics in a representative neck rotation experiment: (a) angular velocity versus time; (b) angular acceleration versus time.

To allow measurement of axonal fiber direction, DTI images of the same subject are acquired using at least 30 non-colinear gradient directions^[41]^, and TORTOISE^[42]^ is used to estimate the diffusion tensor, fractional anisotropy (FA), and labels of major axonal fiber tracts. The diffusion tensor provides the primary direction of axonal fiber alignment at every brain image voxel, allowing computation of axonal fiber strain *E*_*f*_ from the G-L strain tensor. FA estimates the degree of fiber dispersion at a given brain voxel; in this work, only voxels with FA values > 0.2 are considered for computation of *E*_*f*_, which is a standard criterion^[43]^ implemented to focus on regions of high fiber alignment.

## 3. Computational Modeling and Simulation

### 3.1. Constitutive Framework

To capture the nonlinear features of the mechanical response of brain tissue under large strain and dynamic strain rates (e.g., strain stiffening, compression-tension asymmetry, and rate-sensitivity), the viscous dissipation-based visco-hyperelastic modeling framework^[19,44,45]^ is employed in this work. First proposed by Pioletti et al.^[44]^ for modeling short-time (i.e., high strain rate) and large-deformation responses of biological tissues, this framework assumes that the entropy production under dynamic strain rates occurs entirely due to viscous dissipation in the material, which in turn is modeled using the objective strain rate tensor as an external state variable. In its most general form^[45]^ for a compressible and anisotropic material, this constitutive model is written as

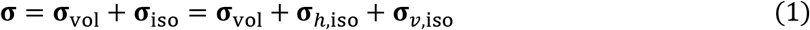

with

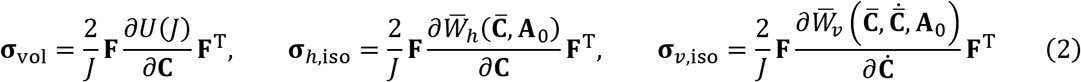

where **σ** is the Cauchy stress tensor, which is additively decomposed into separate volumetric (**σ**_vol_) and isochoric (**σ**_iso_) components. The latter is further split into hyperelastic (**σ**_*h*,iso_) and viscous overstress (**σ**_*υ*,iso_) components. Specifically, the rate-independent volumetric stress **σ**_vol_ is derived from the volumetric energy density *U*, which is a function of *J* = det(**F**) (**F** is the deformation gradient tensor). **C** = **F**^T^**F** is the right Cauchy-Green deformation tensor. Further, the isochoric hyperelastic stress **σ**_*h*,iso_ that is associated with the quasi-static material response is derived from a strain energy density function 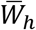, which depends on the modified right Cauchy-Green deformation tensor 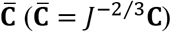 and the structure tensor **A**_0_. Structure tensor accounts for anisotropy in the material response, and is given as **A**_0_ = **a**_0_ ⊗ **a**_0_, where **a**_0_ is the unit vector in the fiber direction. Finally, the viscous overstress **σ**_*υ*,iso_ is associated with the dynamic rate-dependent material response and is derived using a viscous dissipation potential 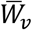, which is a function of 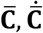, and **A**_0_.

In the present study, the Simo-Miehe form^[46]^ of volumetric energy density is employed,

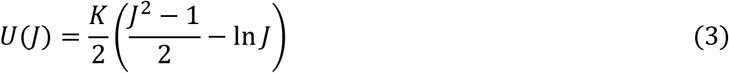

where *K* is the bulk modulus of the material. Equation (3) is especially suited for weakly compressible materials^[45]^ (with relatively large bulk modulus), and has been used to model the volumetric response of brain tissue in several studies^[47,48]^.

Regarding the isochoric stress components in Eq. (2), the present study assumes that the effect of the structural anisotropy of brain tissue on its mechanical response is very small and can be neglected. This is based on recent evidence in the literature^[9,18]^ that suggests that brain white matter exhibits relatively insignificant mechanical anisotropy. The one-term Ogden strain energy density function^[26]^ is utilized to model the quasi-static hyperelastic mechanical response (isotropic),

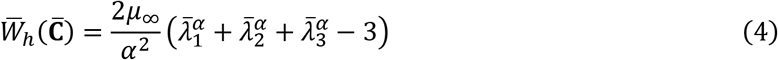

where 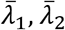, and 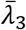 are the distortional principal stretches, *μ*_∞_ is the quasi-static initial shear modulus, and *α* is the compression-tension asymmetry parameter, which captures the nonlinearity in the stress versus strain response in the large deformation regime. The one-term Ogden model is chosen because it has been shown in several recent studies^[14,18,49]^ to be the best performing hyperelastic model for brain tissue, which can capture major characteristics of quasi-static brain tissue deformation: strain stiffening, compression-tension asymmetry, and sharp increase in shear modulus with superimposed compression but not with superimposed tension. In addition, the Ogden model can simultaneously capture multiple homogeneous deformation modes, which is important for accurately predicting the mechanical response under triaxial loading conditions^[50,51]^.

To capture the isochoric viscous overstress, the three-parameter Upadhyay-Subhash-Spearot (USS) viscous dissipation potential^[19,52]^ is employed,

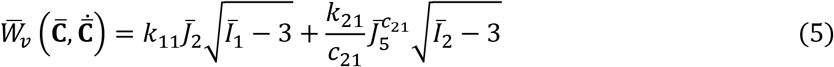

where 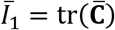 and 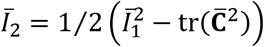 are principal invariants of 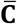, and 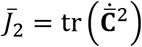 and 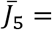 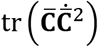 are the scalar invariants of tensors 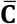 and 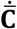. *k*_11_ and *k*_21_ are the linear and nonlinear rate sensitivity control parameters, respectively, which control the rate at which elastic moduli increase with strain rate, and *c*_21_ is the rate sensitivity index, which captures the curvature associated with this increase. The USS model can flexibly capture the nonlinear strain-stiffening and strain rate-stiffening responses that are characteristic of the mechanical response of brain tissue under high strain rate loading^[53]^, and can simultaneously fit multiple deformation modes with good accuracy^[19]^.

Substituting Eqs. (3–5) in Eqs. (1–2), the following constitutive equation is obtained (see derivation in section SM1 of the supplementary material),

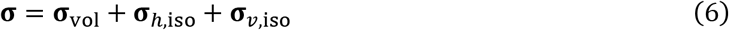

with

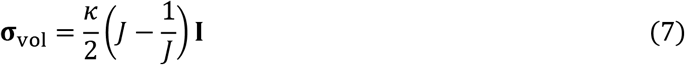

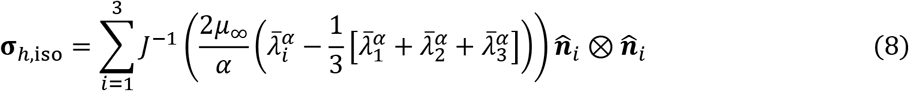

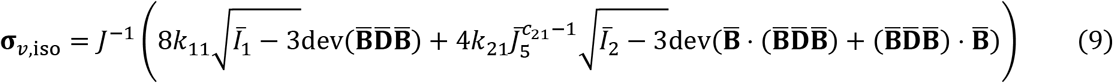

where **I** is the unit symmetric tensor and 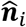 are the principal directions of the modified left Cauchy-Green deformation tensor 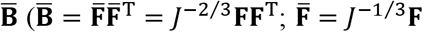 is the modified deformation gradient). Further, 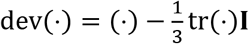 is the Eulerian deviatoric operator, and 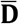 is the modified rate of deformation tensor, which is the symmetric part of the modified spatial velocity gradient 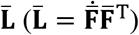.

The constitutive model in Eqs. (6–9), hereinafter referred to as the O-USS model, is utilized in the present study to simulate the response of all parenchymal brain substructures: deep and cortical gray matter, corona radiata, corpus callosum, cerebellar gray and white matters, and brainstem. The subarachnoid space (SAS) is modeled as a generalized Maxwell LVE material with a Prony-series (*N* = 1) representation of the relaxation function (*G*(*t*)),

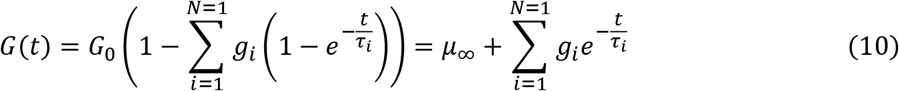

where *G*_0_ and *μ*_∞_ are the short-term and long-term (i.e., quasi-static) shear moduli, respectively, *g*_*i*_ are constant parameters, and *τ*_*i*_ are the relaxation time constants. Note, 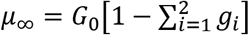. Further, falx and tentorium are modeled as linear elastic materials. Ventricles and cerebrospinal fluid are modeled as a viscous fluid using the shear viscosity (to capture shear behavior) and the Murnaghan-Tait equation of state^[54]^ (to capture hydrostatic pressure):

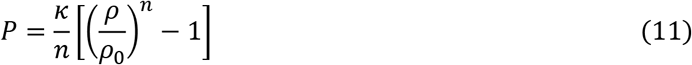

where *P* is the pressure, *ρ* is the density (*ρ*_0_ is density at zero pressure), and *n* is a constant; for fluids similar in composition and viscosity to water (CSF is composed of 99% water^[55]^), *n* = 7.15^[56]^. Finally, the skull is modeled as a rigid linear elastic material.

To highlight the efficacy of the O-USS model in capturing 3D brain dynamics under rotational acceleration loading, it is compared with two commonly employed constitutive equations for head models in the literature: (i) generalized Maxwell LVE model^[13]^, and (ii) Ogden model-based linear visco-hyperelastic (O-LVHE) model^[5,57]^. For the LVE model, a two-branch (*N* = 2) Prony-series representation of the relaxation function (Eq. (10)) is employed. For the O-LVHE model, which assumes an additive decomposition of the total stress into nonlinear hyperelastic and linear viscoelastic components, a reduced form of the relaxation function in Eq. (10), i.e., 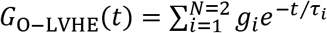, is used. To capture the hyperelastic stress component, the Ogden strain energy density function (Eq. (4)) is used.

### 3.2. In-ex-vivo Hybrid Parametrization

Figure 3 illustrates the O-USS model parametrization procedure proposed in the present study. In the first step of this procedure (Fig. 3(a)), which is detailed in a recent work by the authors^[29]^, the frequency-dependent storage and loss moduli at each MRE brain voxel are used to calibrate a two-branch generalized Maxwell LVE model,

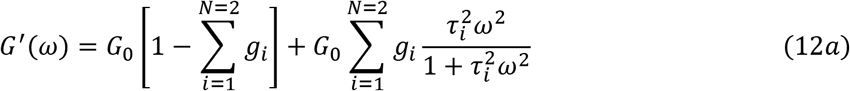

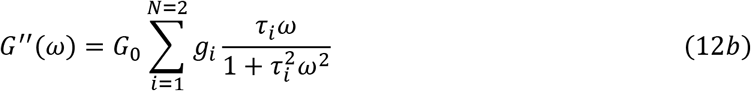

where *G*′ and *G*′′ are the storage and loss moduli, respectively, and *ω* is the applied frequency (30, 50 and 70 Hz), and *g*_*i*_ and *τ*_*i*_ are detailed in Eq. (10).

**Figure 3.**
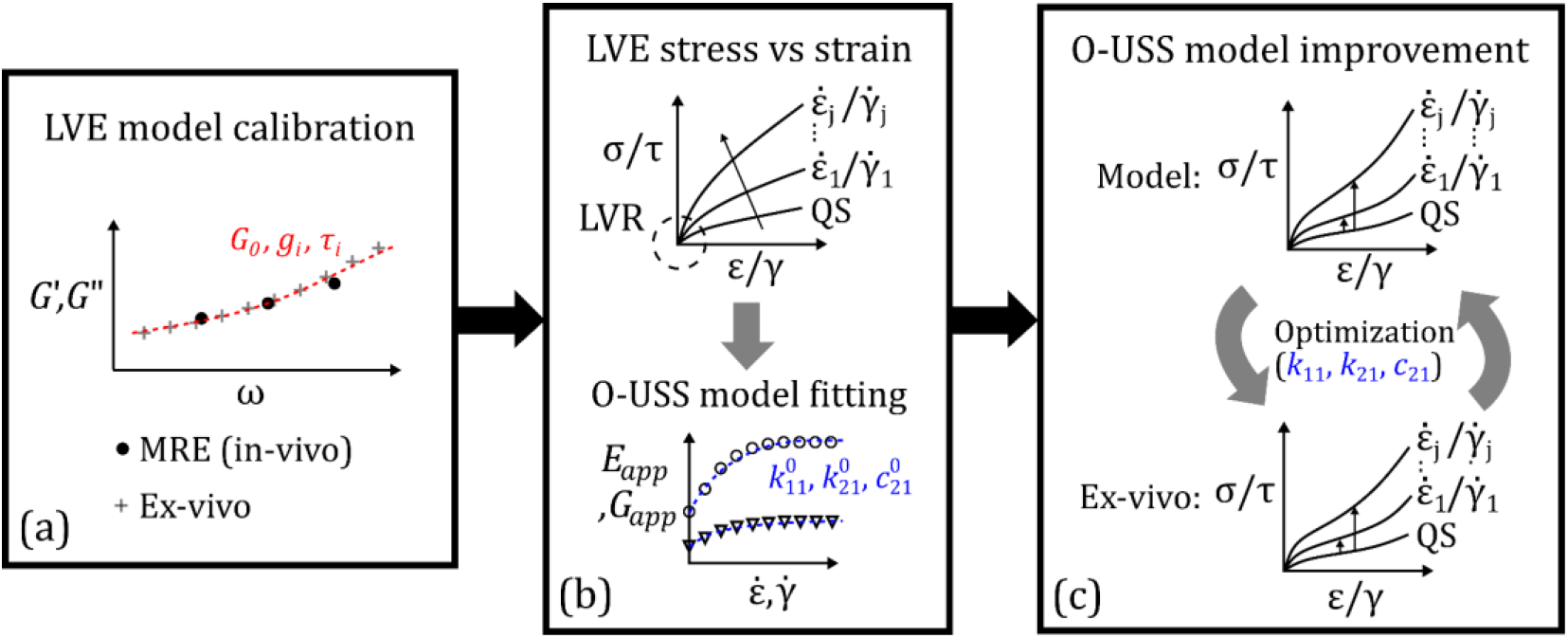
Schematic illustration of the in-ex-vivo hybrid parametrization procedure developed in this study. (a) In the first step, frequency-dependent storage and loss moduli from in-vivo MRE and selected ex-vivo data is fit to the generalized Maxwell LVE model. (b) In the second step, the predicted LVE stress-strain response at multiple constant strain rates is used to obtain the apparent elastic moduli strain rate spectra, which are fit to the O-USS model. (c) In the third step, the O-USS model parameters are refined via a constraint optimization against ex-vivo large deformation stress versus strain data.

As the employable actuation frequency range in in-vivo MRE is limited due to issues related to depth of wave penetration, resolution, noise, and human subject comfort^[39]^, data from an ex-vivo dynamic shear study in the literature with a wider frequency bandwidth is combined with the in-vivo data before model calibration^[29]^. An optimization algorithm is employed to select a set of data that follows similar moduli versus frequency trends as observed in the in-vivo MRE response (details can be found in Alshareef et al.^[29]^). The output of this step is a set of five LVE model parameters (*G*_0_/*μ*_∞_, *g*_1_, *τ*_1_, *g*_2_, and *τ*_2_) for each voxel in the brain; average values of these parameters for different substructures for the three subjects are provided in the supplementary material.

With the LVE model parameters from the first step, a recently proposed frequency to strain rate domain conversion technique^[58–60]^ is utilized to obtain the O-USS visco-hyperelastic model parameters (Fig. 3(b)). In this method, the LVE properties are first used to predict the stress versus strain responses under shear and uniaxial deformation modes, at multiple constant strain rates. A 0.001-200 s^−1^ strain rate range is chosen, which is the approximate range of strain rates in real-life brain injury events^[61]^. The stress-strain responses are then used to compute the apparent elastic moduli-strain rate spectra^[58]^ at each brain voxel,

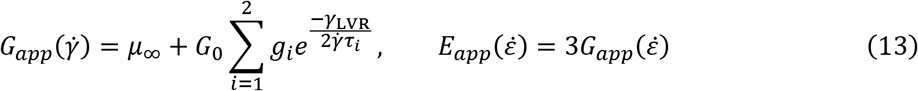

where *G*_*app*_ and *E*_*app*_ are strain rate dependent apparent shear and uniaxial moduli, respectively; *γ* and *ε* denote shear strain and uniaxial strain, respectively. *γ*_LVR_ is the linear viscoelastic regime (LVR) limit, which is approximately 0.01 mm/mm for brain tissue^[22–24]^. Equation (13) represents the approximate average shear and uniaxial moduli of a material in its LVR when the rheological functions are independent of the input strain level. Note that this equation assumes incompressibility, which is a reasonable assumption for brain tissue^[49]^. The apparent elastic moduli-strain rate spectra are fit using the rate-sensitive tangent moduli equations of the O-USS model^[19]^,

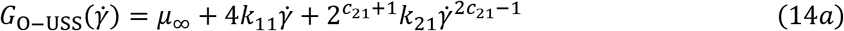

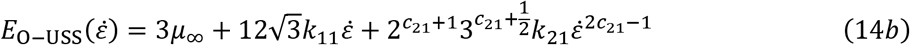

This yields a set of optimized viscous dissipation potential parameters: 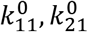, and 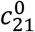, for each brain voxel. Combined with *μ*_∞_ obtained in the previous step, this second step yields four visco-hyperelastic model parameters for each voxel in the brain.

Finally, in the third step, the ex-vivo large deformation stress versus strain data of different brain regions from the literature are utilized to inform the visco-hyperelastic model about the large-strain regime of brain tissue deformation. Four criteria are implemented for the selection of ex-vivo studies for this purpose: the stress-strain data should be (a) under large deformations (≥ 20% engineering strain), (b) under all three primary deformation modes of compression, tension, and shear, (c) at both low and high strain rates, and (d) ideally be from human brain tissue (not from an animal model). The criterion (b) is critical in the development of 3D computational models, where model parameters calibrated from a single deformation mode can result in thermodynamic instabilities and poor accuracy when simulating a general triaxial deformation mode^[19,50,51]^. A summary of the selected ex-vivo studies for all brain substructures is provided in the supplementary material. Out of the seven brain substructures, ex-vivo data of four regions (cortical gray matter, deep gray matter, corona radiata, and corpus callosum) fulfill all the four selection criteria, whereas for brain stem and cerebellar gray and white matter tissues, all but (d) criteria are fulfilled (stress-strain data is from porcine brain).

With the selected ex-vivo data sets, the compression-tension asymmetry parameter (*α*) for each brain region is first obtained by fitting the experimental quasi-static (taken as the smallest investigated strain rate) stress versus strain response under the three primary deformation modes with the stress-strain equations of the Ogden model, which is the quasi-static component of the O-USS model (details of the hyperelastic model fitting procedure is provided elsewhere^[50]^). This finalizes the first estimate of the visco-hyperelastic model parameters: *μ*_∞_, *α*, 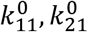, and 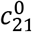. Next, a nonlinear constrained optimization problem is employed to obtain “refined” viscous dissipation model parameters (see Appendix A). This optimization minimizes the relative difference in viscous overstress (total dynamic stress minus quasi-static stress) between ex-vivo experimental data and model predictions while at the same time minimally affecting the previously obtained estimate of apparent elastic moduli strain-rate spectra from the in-vivo MRE data. In this way, the set of optimized voxel-wise O-USS model parameters obtained in this step (Fig. 3(c)) result in a constitutive model that is calibrated to capture the subject-specific in-vivo and strain rate dependent small-strain response of brain (i.e., linear viscoelastic), and at the same time is informed about the large strain response features (nonlinearity, compression-tension asymmetry, and rate sensitivity) with the help of available ex-vivo data in the literature. Finally, the bulk modulus (*κ*) for all brain tissue types is assumed to be 2.19 GPa in this study, which is a value widely used in the literature^[13,62]^. The complete set of regional average O-USS model parameters for a particular subject, S01, are listed in Table 1, along with the material properties of finer brain substructures obtained from ex-vivo studies in the literature. For the calibrated O-USS model parameters of the other two subjects, see the supplementary material.

**Table 1.**
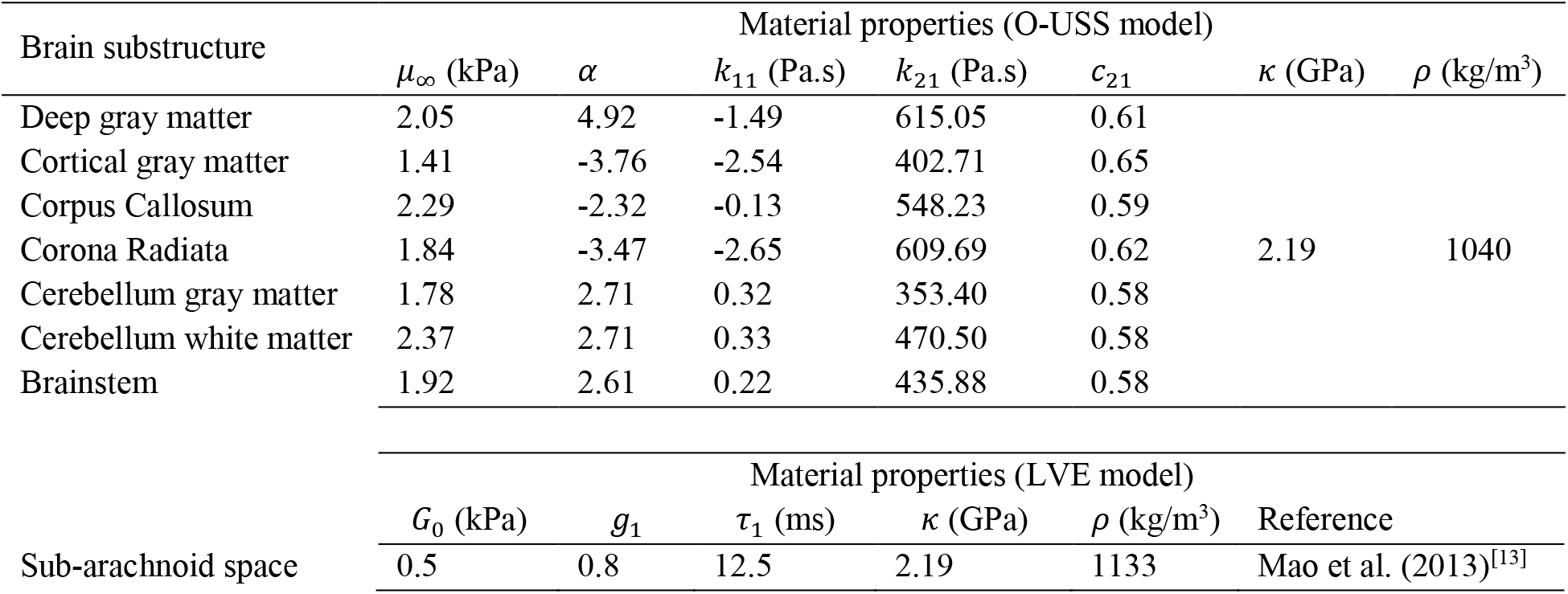
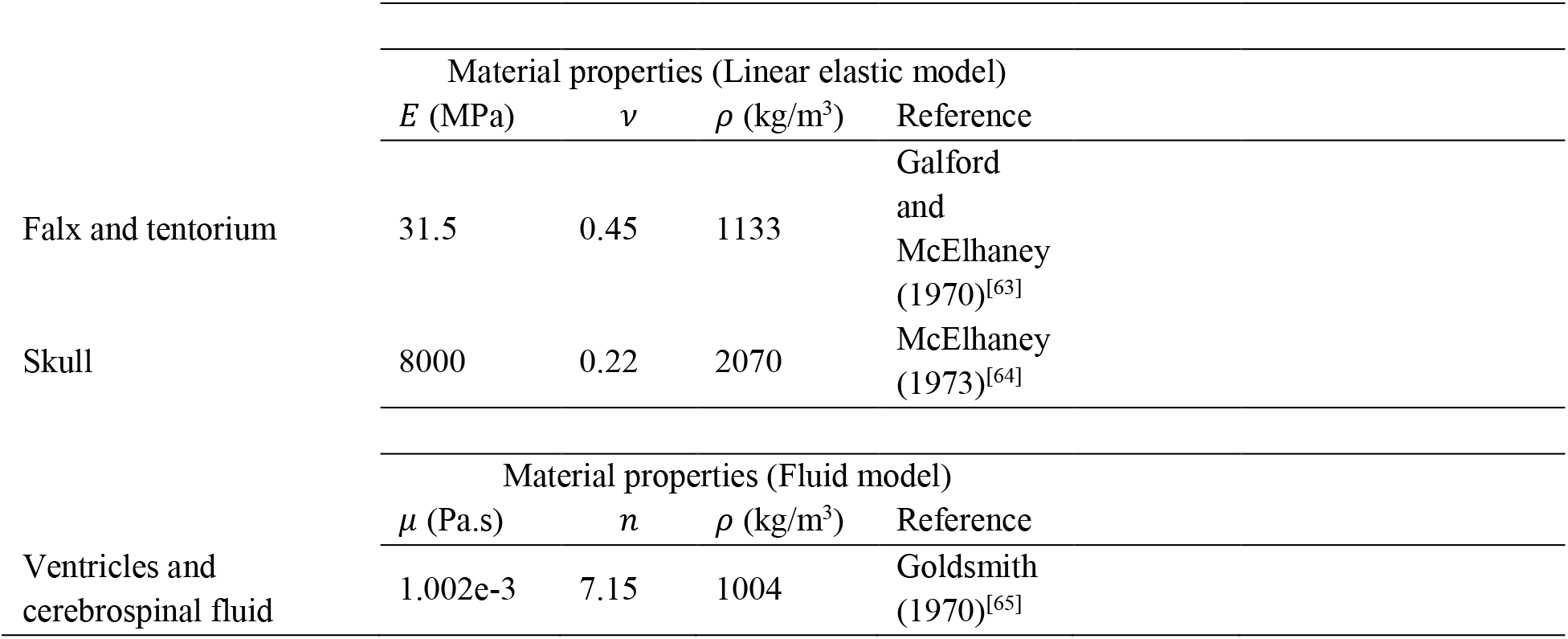
Material properties of the various brain substructures (for subject S01), used in this study.

### 3.3. Computational Approach

The Material Point Method (MPM)^[66,67]^ is used in this study to numerically simulate the full-field dynamic response of brain. MPM is a meshless, mixed Lagrangian-Eulerian method, in which the material continuum is partitioned into a collection of particles that define the Lagrangian state of the system, while the equations of motion are solved on a fixed background Eulerian grid. This mixed formulation offers several significant advantages when it comes to brain/head models^[6,47,62]^: (i) voxelated biomedical images can be directly imported as material points, thus reducing the computational cost of model generation; (ii) no issue of mesh distortion that is especially prominent in finite element models undergoing large deformations; (iii) robustness in simulations involving multi-phase systems such as brain; (iv) efficient handling of large bulk-to-shear modulus ratios without volumetric locking artifacts. Details of the implementation of the O-USS model-based head simulations in MPM are provided in the supplementary material. For the subject S01, material properties in Table 1 and boundary conditions in Fig. 2 are employed.

Note that with LVE- and O-LVHE models, MPM simulations led to stress inaccuracies, which manifests as a checker-boarding pattern in the simulated stress/strain-field (see supplementary material): this is a known limitation of MPM, which is exacerbated in the case of advanced history-dependent constitutive models and at large deformations^[68,69]^ (LVE and O-LVHE models predict much larger deformations compared to the USS model). Thus, for head models with these constitutive models, finite element analysis is employed (details in Alshareef et al.^[29]^). For subject S01, the LVE model parameters of the LVE and O- LVHE models (obtained from first step of parameterization) are provided in the supplementary material. Parameter *α* for the O-LVHE model is taken from Table 1; for both models, boundary condition in Fig. 2 is applied.

## 4. Quantitative Model Validation Framework

In this study, a robust quantitative validation method is proposed that compares full-field time-varying tensorial and scalar strain measures obtained from the simulation and the corresponding in-vivo tMRI imaging. For the tensor-components of the G-L strain, the volume fraction of brain that has strain within a specific range (say, *m* to *n*) is first analyzed,

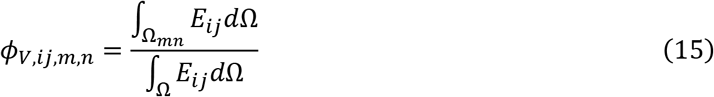

Here, *E*_*ij*_ is a component of the G-L strain tensor, **E**, and Ω_*mn*_ is the subset of the overall domain Ω where *m* ≤ *E*_*ij*_ ≤ *n* holds at a given time. Equation (15) results in time-varying scalar values of volume fraction for each “bin” of a strain component, *ϕ*_*V*,*ij*,*m*,*n*_. These values from simulations and experimental results for a given time step can then be compared to assess the accuracy of the model in predicting the strain tensor field. In this study, only the time step that corresponds to maximum experimental strain values, when the brain is most vulnerable to injury, is considered.

Even though useful, the volume fraction-based metric does not capture the actual overlap of predicted volumes with the corresponding experimental observations. Thus, the Sørensen-Dice coefficient (*D*) is employed to assess the spatial accuracy of the computational model. For a given experimental region of interest, *V*_*exp*_, and the corresponding model prediction, *V*_*mod*_,

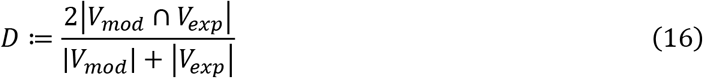

where ∩ denotes intersection of two regions. The *D* score ranges from 0 (i.e., no overlap) to 1 (i.e., a perfect overlap). For G-L strain tensor components (*E*_*ij*_), *D* is computed for each of the “bins” (*m* ≤ *E*_*ij*_ ≤ *n*). The Sørensen-Dice coefficient is the primary spatial validation metric employed in this study. In addition to the G-L strain, spatial accuracy of the prediction of two time-independent scalar strain measures are also computed: (i) maximum axonal strain (MAS), i.e., the maximum of axonal fiber strain (*E*_*f*_) over the entire loading duration at each brain voxel, and (ii) cumulative maximum principal strain (CMPS), which is the corresponding maximum of maximum principal strain (MPS) for a given voxel. CMPS and MAS are commonly employed for quantifying injury in the brain biomechanics community^[70]^. For example, the “injured” volume fraction of brain following an impulse loading can be computed based on CMPS- or MAS-based injury thresholds^[57,61,70]^. In this study, *D* score is computed for the predicted volume fraction of brain where CMPS and MAS are above certain thresholds.

The CORrelation and Analysis objective rating^[71]^ (CORA) and correlation score^[72]^ (CS) are employed in this study as temporal validation metrics. CORA quantifies agreement between two time-varying signals (or responses) in terms of their phase, size, and progression; CORA scores range from 0 (poor agreement) to 1 (perfect agreement). Here, the cross-correlation method for calculating CORA is used, with the interval of evaluation is manually set to [0 ms, 117 ms]. This time-range comprises of the first seven datapoints from tMRI and features both the first major and the second minor strain peaks (see Section 5). All other settings are adopted from the “recommended CORA parameters” for head models^[73]^. For the time-varying full-field prediction of G-L strain components, MPS, and *E*_*f*_, voxel-wise CORA values are averaged across the brain using experimental peak strain values at each voxel as weighting factors. Thus, voxels with a higher strain magnitude, which are more relevant in brain injury prediction, contribute more to the CORA score than those that have a smaller strain. Further, an un-weighted CORA score is reported for the time-varying peak (evaluated at 95- percentile level) and average strains (evaluated at 50-percentile level).

CS is derived from the normalized integral square error (NISE), which compares a pair of time-varying responses in three aspects: phase shift, amplitude difference, and shape difference. CS values range from 0 (poor agreement) to 100 (perfect agreement). The [0 ms, 117 ms] interval is used for evaluation. Like CORA, weighted CS is reported for the time-varying full-field predictions of G-L strain components, MPS, and *E*_*f*_. For the time-varying peak and average strains, an un-weighted CS is reported.

Lastly, the difference between predicted and experimentally observed peak and average strains at the time step corresponding to maximum strain from tMRI is reported in the form of an absolute relative error (with respect to the experimental strain value).

## 5. Full-field Strain Response and Model Performance

### 5.1. Qualitative comparison between experimental and simulated response

During a neck rotation experiment, the skull rapidly decelerates from an initially constant angular velocity (Fig. 2), which causes the internal brain substructures to deform and develop strain. Figure 4 compares the time evolution of the three axial in-plane G-L strain components (*E*_*xx*_, *E*_*yy*_ and *E*_*xy*_) from simulation with the experiment data of a particular subject, S01, at four equally spaced axial brain slices. Time-independent MAS and CMPS fields are also shown. The time steps (9 ms–63 ms) are the first four time-instants when experimental data is available. The column representing the first frame in tMRI at time *t* = 9 ms is common for each of the three strain components, which are all nearly zero at that time. Note that the slice *z* = 0 mm corresponds to an axial brain layer passing through the genu of the corpus callosum.

**Figure 4.**
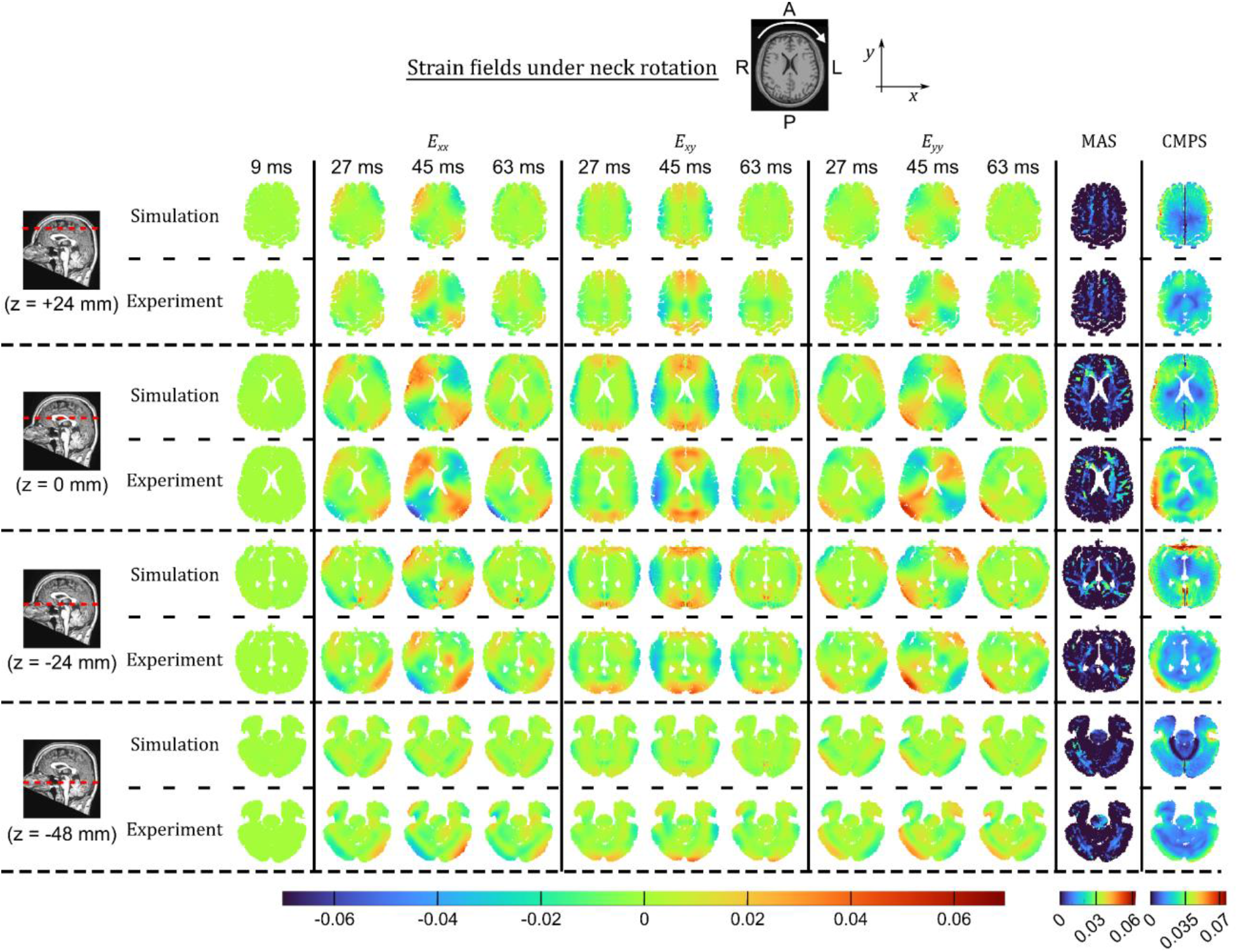
Comparison of the evolution of in-plane Green-Lagrange strain fields, and the MAS and CMPS strain fields, from head model simulations and tMRI of neck rotation experiment, at four axial brain slices. Each column in the case of the Green-Lagrange strain corresponds to a specific experimental time-step.

From Fig. 4, both the simulated and experimental G-L strain-fields reach maxima at approximately 45 ms, which roughly corresponds to the time of maximum negative head velocity during the neck rotation experiment (Fig. 2). At both 45 ms and its preceding time-step, a good qualitative agreement is observed between simulated and experimental strain-fields, both in terms of their spatial distribution and the actual range of strain values. While an anti-symmetric spatial strain distribution of negative (i.e., compressive) and positive (i.e., tensile) strains roughly about the center of the brain slices is observed in the case of the normal strain components (*E*_*xx*_ and *E*_*yy*_), a symmetric spatial distribution is observed for the in-plane shear strain. At 63 ms, the strain is much lower, and the correlation between the spatial strain distribution between simulation and experiments is weak at the periphery of the brain slices (near the brain-SAS interface). After 63 ms, the strains relax following a minor peak; these strain fields are not shown. The MAS and CMPS fields from tMRI are also in a good visual agreement with the corresponding simulation results.

Regardless of the strain type, it is seen that the outer parts of brain are exposed to higher strains, which concurs with the observation in a previous work that noticed that the shear waves resulting from rotational deceleration decay while traveling from the outer to the inner brain regions^[6]^. Relatively large MAS are observed in the left-frontal and right-posterior corona radiata, suggesting a contrecoup pattern. Finally, even though very small compared to the in-plane components, the out-of-plane strain components (*E*_*xz*_, *E*_*yz*_ and *E*_*zz*_) predicted by the model also show a reasonable spatial agreement with the corresponding experimental strains (see supplementary material).

### 5.2. Quantitative model accuracy and validation

To quantitatively assess the accuracy of the head model, the present work employs three types of metrics: spatial validation metrics, temporal validation metrics, and relative errors of prediction of strain maxima. Figures 5(a-c) show bar graphs of the volume fractions of brain under different in-plane G-L strain ranges at 45 ms (bin width is 0.015 mm/mm), from simulation and experiment. For each of the three strain components, the brain volume fraction under a positive strain range is approximately equal to the volume fraction under the corresponding negative strain range; this is a direct consequence of the symmetric/anti-symmetric strain-fields observed in Fig. 4. Overall, a good agreement between the simulation and experiment is observed, both in terms of the volume fraction under individual strain range bins, and the overall range of values of individual strain components.

**Figure 5.**
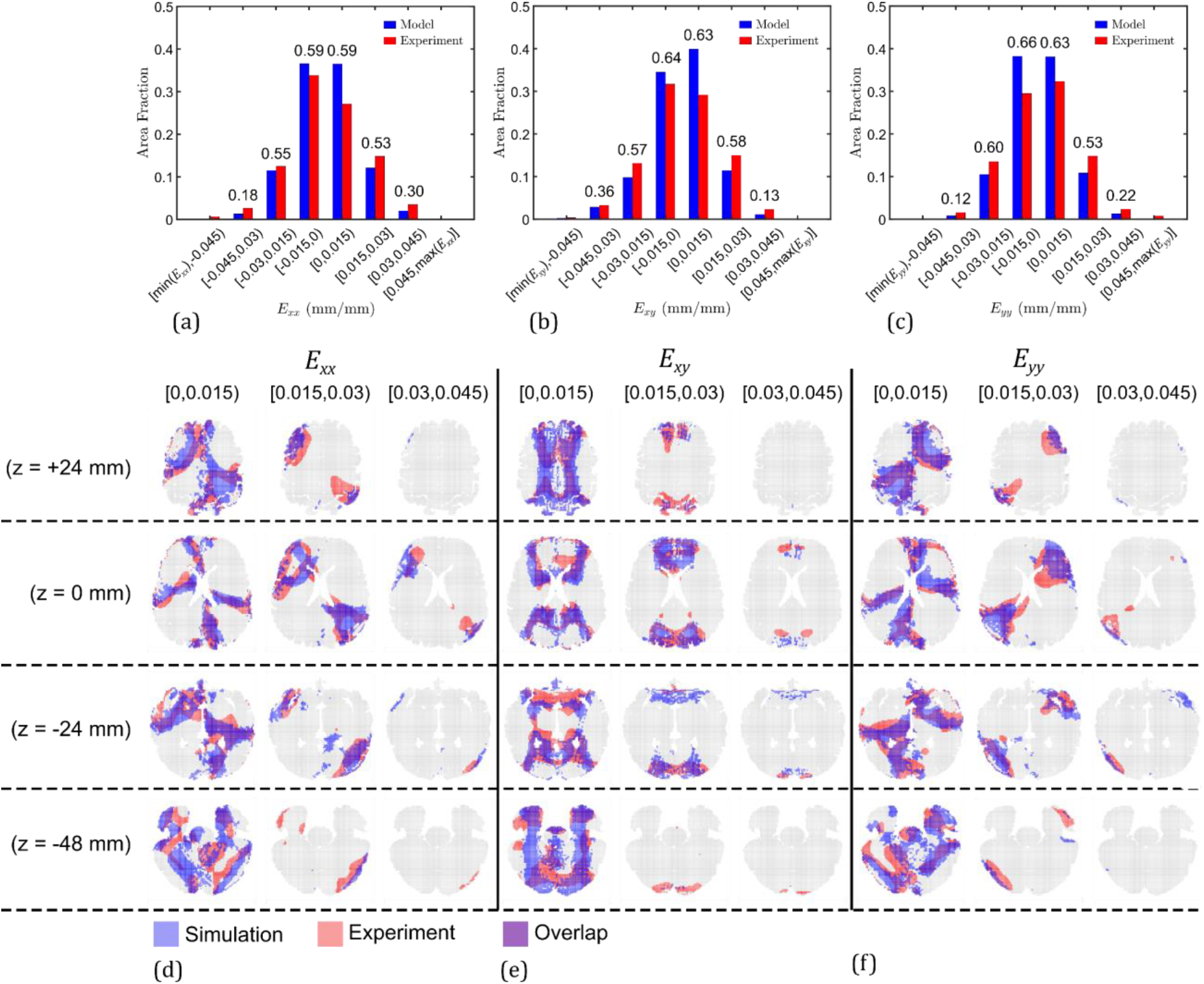
Bar graphs showing volume fraction of brain exposed to different ranges (or bins) of Green-Lagrange strains at 45 ms (corresponding to highest strain in experiment): (a) *E*_*xx*_, (b) *E*_*xy*_, and (c) *E*_*yy*_. Values written over individual bars are the Sørensen-Dice coefficients of the overlap between simulated and experimental brain volumes under that particular strain range. Brain regions under three strain bins (i.e., [0, 0.015), [0.015, 0.03), and [0.03, 0.045)) at 45ms time step, shown in four axial slices, highlighting the overlap (purple) in the predicted (blue) and experimentally observed (red) strain regions: (d) *E*_*xx*_, (e) *E*_*xy*_, and (f) *E*_*yy*_.

To further analyze the spatial agreement of the in-plane G-L strain components, the Sørensen-Dice coefficient (*D*) metric is indicated over individual bars in Figs. 5(a-c). For the four bins in the −0.03 mm to 0.03 mm range that cover more than 95% of the brain by volume, *D* is always greater than 0.5, showing a significant overlap between the predicted and observed volumes, as visualized in Figures 5(d-f). Even for the [0.03, 0.045) bin that corresponds to a very small brain volume, there is a general agreement in the locations of these high strain regions between the simulation and the experiment, though the *D* overlap scores for this bin are lower owing in part to its smaller size.

Table 2 lists the *D* scores averaged across the six bins in the −0.045 mm to 0.045 mm range (covering more than 99% brain volume), for all the six G-L strain components. Note that superscript 0 with individual strain components denote their values at the time-step corresponding to peak strains, which is 45 ms in this case. For the out-of-plane strain components, the *D* score is approximately half of the score for in-plane strain components. This reduction in spatial agreement is attributed to the high uncertainty in experimental measurement in the out-of-plane z-direction (I/S), along with a low signal-to-noise ratio (SNR) due to small strain magnitudes.

**Table 2.** Model validation statistics.

| Spatial Validation |  | Temporal Validation |  |  | Peak and Average Strain Errors |  |  |
| --- | --- | --- | --- | --- | --- | --- | --- |
| Strain | Average<br>Sørensen-Dice<br>Coefficient<sup>a</sup> ($\bar{D}$) | Strain | Weighted<br>average CORA<br>score | Weighted<br>average CS<br>score | Strain | Peak<br>strain<br>error (%) | Average<br>strain error<br>(%) |
| $E_{xx}^0$ | 0.46 | $E_{xx}$ | 0.43 | 49.60 | $E_{xx}$ | 6.55 | 30.10 |
| $E_{xy}^0$ | 0.49 | $E_{xy}$ | 0.44 | 51.95 | $E_{xy}$ | 6.50 | 21.21 |
| $E_{yy}^0$ | 0.46 | $E_{yy}$ | 0.47 | 54.73 | $E_{yy}$ | 10.59 | 29.00 |
| $E_{xz}^0$ | 0.24 | $E_{xz}$ | 0.31 | 30.79 | $E_{xz}$ | 11.65 | 16.54 |
| $E_{yz}^0$ | 0.24 | $E_{yz}$ | 0.37 | 35.60 | $E_{yz}$ | 41.07 | 13.15 |
| $E_{zz}^0$ | 0.21 | $E_{zz}$ | 0.27 | 21.29 | $E_{zz}$ | 48.96 | 37.35 |
| **MAS** | 0.47 | $E_f$ | 0.42 | 44.52 | $E_f$ | 3.50 | 6.34 |
| **CMPS** | 0.44 | MPS | 0.71 | 83.16 | MPS | 4.31 | 2.87 |

<sup>a</sup>For Green-Lagrange strains, averaging across six strain bins in the -0.045 to 0.045 mm/mm range; for MAS and CMPS, averaging from 50- to 95-percentile MAS and CMPS thresholds, respectively.

Further, the spatial overlap of brain volumes between simulation and experiment where MAS and CMPS are above a certain threshold are analyzed in Fig. 6. Figs. 6(a,c) plot *D* score as a function of threshold MAS and CMPS, respectively. Not surprisingly, the *D* values start from a maximum value of ~1 when both simulation and experiment show the entire brain volume as above the zero-threshold, and decrease thereafter for larger threshold values as the predicted volumes become smaller. Figure 6(b) shows the overlap between the predicted and observed brain volumes with MAS greater than its 50- and 95- percentile thresholds (MAS50 and MAS95). These two thresholds are often considered as average and peak MAS, respectively^[27,41]^. A good level of spatial agreement is evident from these maps. Notably, the head model accurately predicts the regions of highly strained axonal fibers (> MAS95), which are visible as scattered “islands” (marked by dashed circles) on the axial slice image. Overall, the average *D* score for thresholds from 50- to 95-percentile is 0.47 (see Table 2).

**Figure 6.**
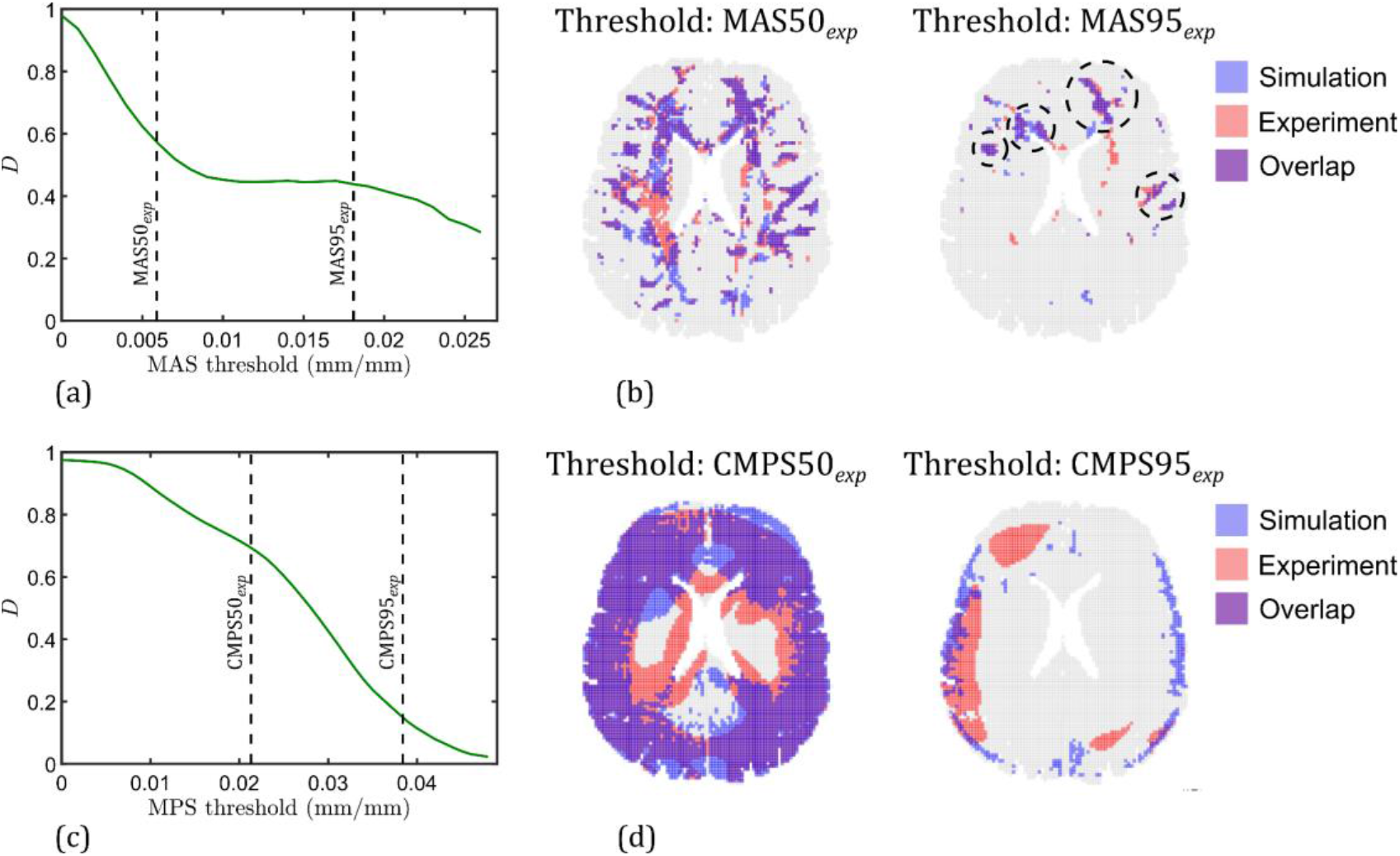
Sørensen-Dice coefficient as a function of threshold strain, for (a) maximum axonal strain (MAS), and (c) cumulative maximum principal strain (CMPS). Comparison of the predicted and experimentally observed regions of brain (slice: z = 0 mm) where (b) MAS is greater than its 50- and 95-percentile thresholds, and where (d) CMPS is greater than its 50- and 95-percentile thresholds.

Unlike the case of MAS where *D* was consistent from 50- to 95-percentile thresholds, a sharp decay in *D* score is noticed for CMPS. This time, the average *D* score for thresholds from 50- to 95-percentile is 0.44 (Table 2). Examining the regions of overlap between experiment and model in Fig. 6(d), a greater disagreement is observed in the case of the CMPS95 threshold: while the simulation predicts highly strained regions located symmetrically on the left and right periphery of the brain, the experimental observations show high CMPS on a larger area that is mainly located only on the right half of the brain. A potential reason for this discrepancy might be that while the computational head model assumes a single set of material properties for each of the different brain substructures that are nearly symmetrically on the left and right sides of brain (leading to a symmetry in the mechanical response as well), in reality, differences in material properties between the left and right sides of the brain may lead to asymmetry in the mechanical response.

Figure 7 plots the evolution of average and peak strains versus time, for the three in-plane G-L strain components, *E*_*f*_, and MPS. Note that for the G-L strain components and *E*_*f*_, which hold both positive and negative values, average (positive) values are evaluated at 75-percentile (due to their symmetric positive and negative values under rotational loading, their 50-percentile value is approximately zero). For MPS, which is strictly positive, average is evaluated at 50-percentile. Peak strain is always evaluated at 95-percentile.

**Figure 7.**
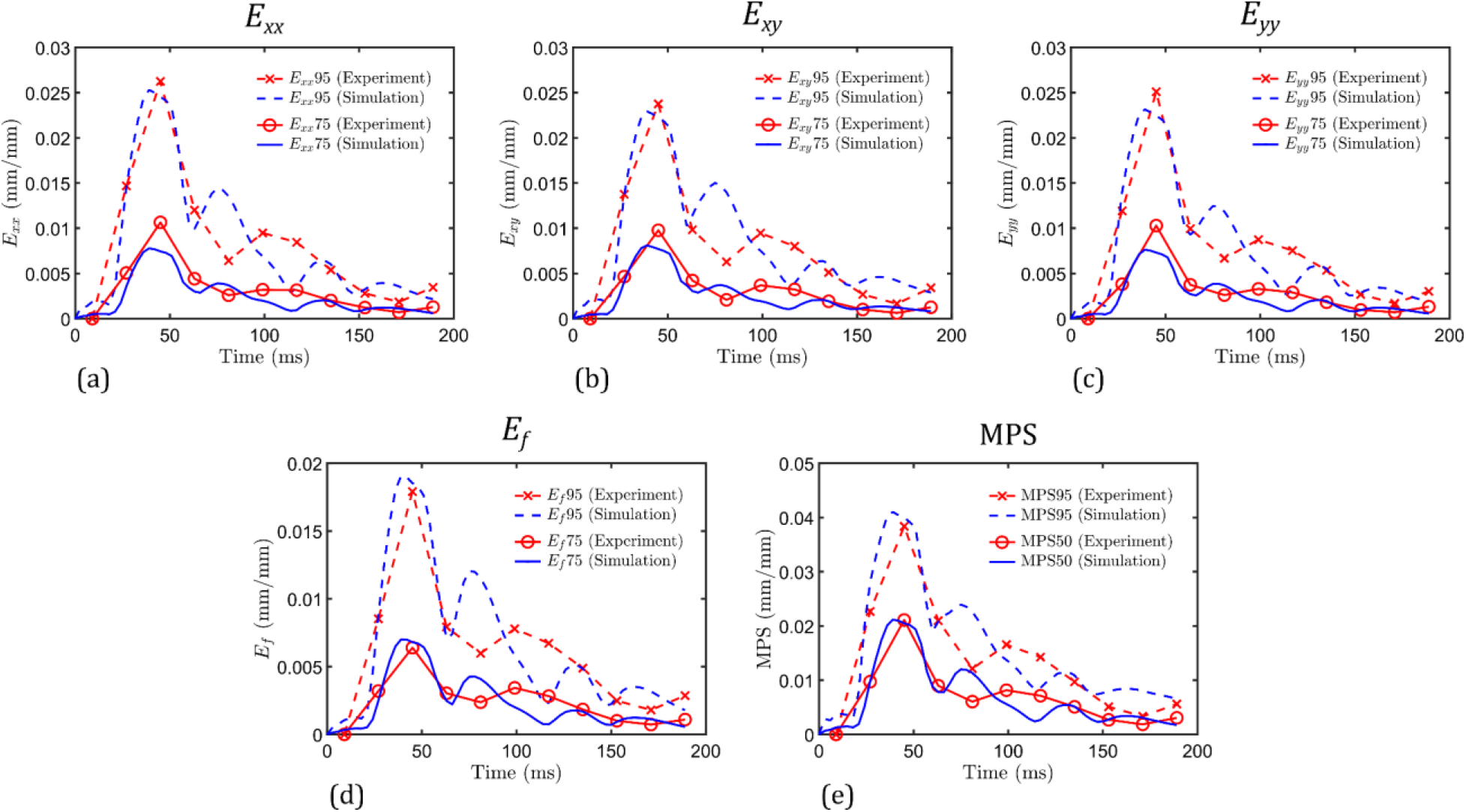
Evolution of the average and peak strains versus time: (a) *E*_*xx*_, (b) *E*_*xy*_, (c) *E*_*yy*_, (d) *E*_*f*_, and (e) MPS.

From Figs. 7(a-e), there is an excellent agreement in the time-evolution of peak and average strains between simulation and experiment, both in terms of the location and the magnitude of the global strain maximum. Specifically, the difference between the time corresponding to global strain maximum from simulation and experiment is approximately 6 ms, which is smaller than the temporal resolution of the experiment that is 18 ms. Table 2 lists the relative error in the prediction of strain at the time-step corresponding to global maximum in the experiment (i.e., 45 ms), for the different strain types. For the in-plane strain components, the relative error of the prediction of peak strain maximum is smaller than the corresponding errors in the average strain maximum. No such correlation exists for the out-of-plane strain components. Between the G-L strain components and the two scalar strain measures (*E*_*f*_ and MPS), the latter are associated with lower relative errors.

After their first maximum, the strain versus time responses in Fig. 7 begin to relax following a second local maximum, though the time this occurs differs between simulation and experiment by approximately 24 ms. Nevertheless, the overall dynamics in terms of the rate of strain decay show good agreement, with the strain magnitudes at the last tMRI frame of 189 ms from simulation and experiment being nearly identical (< 0.01% strain difference). The weighted CORA and CS scores of the temporal agreement of the G-L strain components, *E*_*f*_, and MPS are listed in Table 2. Among the investigated strains, MPS is associated with the highest weighted average CORA and CS scores. Note that these “average” scores evaluate the temporal evolution of strain in every brain voxel. The corresponding un-weighted CORA and CS scores for the average and peak strain versus time responses in Fig. 7 are listed in Table 3. These scores are significantly higher than the average whole-brain scores in Table 2. Further, unlike the whole-brain average scores where the in-plane G-L strain components clearly out-perform the out-of-plane strain components, in terms of the un-weighted CORA/CS scores, there is no such difference between in- and out-of-plane strain components. This shows that the evolution of average and peak values of the entire G-L strain tensor is accurately captured by the head model.

**Table 3.** Un-weighted temporal validation scores for average and peak strain values.

| Strain | CORA score |  | CS score |  |
| --- | --- | --- | --- | --- |
|  | Average strain | Peak strain | Average strain | Peak strain |
| $E_{xx}$ | 0.80 | 0.92 | 92.20 | 96.11 |
| $E_{xy}$ | 0.83 | 0.91 | 94.23 | 95.60 |
| $E_{yy}$ | 0.80 | 0.93 | 92.93 | 96.72 |
| $E_{xz}$ | 0.87 | 0.89 | 93.50 | 95.60 |
| $E_{yz}$ | 0.94 | 0.83 | 96.63 | 94.08 |
| $E_{zz}$ | 0.80 | 0.89 | 90.75 | 94.75 |
| $E_f$ | 0.89 | 0.91 | 93.84 | 95.53 |
| MPS | 0.93 | 0.90 | 96.22 | 96.80 |

## 6. Comparison with Commonly Used Constitutive Frameworks in Head Models

In this section, the simulation results of three subject-specific head models of subject S01, employing the same head geometry and boundary condition but different constitutive models: O-USS, LVE, and O-LVHE models, are compared. Figures 8(a) and 8(b) show the time evolution of peak *E*_*f*_ and MPS (95-percentile level), respectively, for these models. All three models predict a similar location of the first maximum in these strain-time responses. The LVE- and O-LVHE-based models, however, significantly overpredict the magnitude of the maximum strain. Similar observation was made for all other strain types considered in this work at both average (50-percentile) and peak strain levels, resulting in high relative errors as listed in Table 4. Prediction of larger strains by LVE- and O-LVHE-based models can be attributed to the strain-softening behavior in the early part of their stress-strain responses^[74]^, which is the result of a monotonically decreasing stress relaxation function (Eq. (10)). Similarly, the second strain maxima in LVE and O-LVHE models is also significantly larger than that seen in either the O-USS model or experimental observation. Overall, unlike the experimentally observed brain dynamics that feature a gradually decaying strain following the first maximum, the LVE and O-LVHE models predict a strongly oscillatory decay. This results in lower temporal validation scores of these models compared to the O-USS model for both the in-plane Green-Lagrange strains and the two scalar strain measures, *E*_*f*_ and MPS (see Table 4). Note that only the weighted average CORA scores are reported for brevity (CORA and CS scores are positively correlated, as observed from Tables 2 and 3). For the smaller out-of-plane strains, there is no clear difference between the three models in terms of the CORA score.

**Table 4.** Model validation statistics comparison between different constitutive models.

| Spatial Validation |  |  |  | Temporal Validation |  |  |  | Peak Strain Error |  |  |  |
| --- | --- | --- | --- | --- | --- | --- | --- | --- | --- | --- | --- |
| Strain | Average Dice Coefficient<sup>a</sup> ($\bar{D}$) | | | Strain | Weighted average CORA score | | | Strain | Peak strain error (%) | | |
|  | O-USS | LVE | O-LVHE |  | O-USS | LVE | O-LVHE |  | O-USS | LVE | O-LVHE |
| $E_{xx}^0$ | 0.46 | 0.44 | 0.44 | $E_{xx}$ | 0.43 | 0.32 | 0.33 | $E_{xx}$ | 6.55 | 16.99 | 22.25 |
| $E_{xy}^0$ | 0.49 | 0.47 | 0.47 | $E_{xy}$ | 0.44 | 0.30 | 0.31 | $E_{xy}$ | 6.50 | 14.98 | 20.79 |
| $E_{yy}^0$ | 0.46 | 0.50 | 0.50 | $E_{yy}$ | 0.47 | 0.32 | 0.33 | $E_{yy}$ | 10.59 | 11.83 | 16.93 |
| $E_{xz}^0$ | 0.24 | 0.22 | 0.22 | $E_{xz}$ | 0.31 | 0.30 | 0.31 | $E_{xz}$ | 11.65 | 45.87 | 45.87 |
| $E_{yz}^0$ | 0.24 | 0.23 | 0.22 | $E_{yz}$ | 0.37 | 0.31 | 0.31 | $E_{yz}$ | 41.07 | 83.87 | 84.65 |
| $E_{zz}^0$ | 0.21 | 0.20 | 0.20 | $E_{zz}$ | 0.27 | 0.28 | 0.29 | $E_{zz}$ | 48.96 | 66.62 | 78.93 |
| **MAS** | 0.47 | 0.38 | 0.38 | $E_f$ | 0.42 | 0.24 | 0.24 | $E_f$ | 3.50 | 39.75 | 42.77 |
| **CMPS** | 0.44 | 0.47 | 0.46 | **MPS** | 0.71 | 0.59 | 0.59 | **MPS** | 4.31 | 26.45 | 33.18 |

<sup>a</sup>Averaging over the six bins for Green-Lagrange strains, and from 50- to 95-percentile thresholds for MAS and CMPS, respectively.

**Figure 8.**
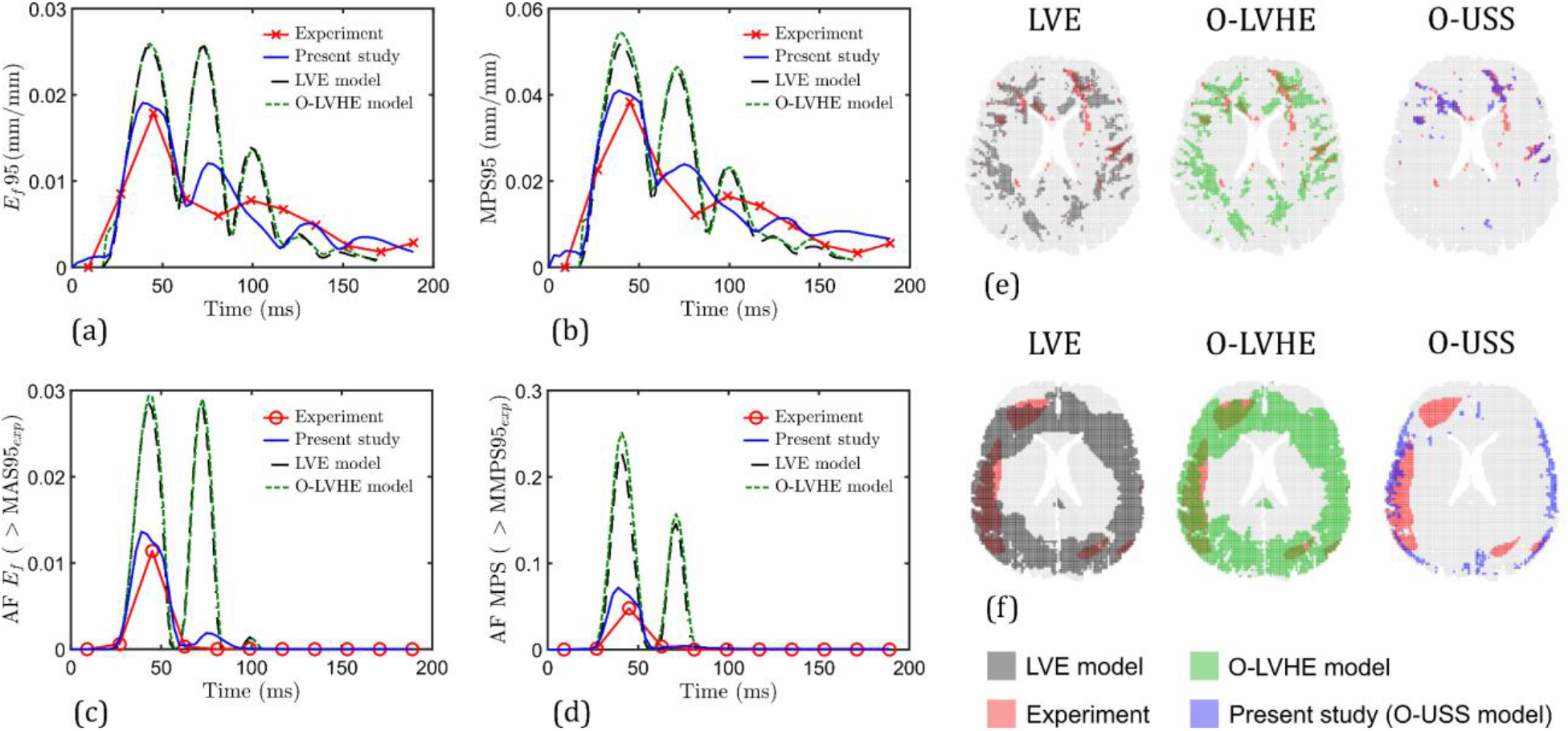
Comparison of the simulated strain-time response of brain resulted by three constitutive models, O-USS model (present study), LVE model, and O-LVHE model: (a) 95-percentile *E*_*f*_ versus time, (b) 95-percentile MPS versus time, (c) volume fraction of brain with *E*_*f*_ greater than 95-percentile MAS (from experiment), and (d) volume fraction of brain with MPS greater than 95-percentile CMPS (from experiment). Predicted and experimentally observed region of brain at z = 0 mm slice where (e) MAS is greater than its 95-percentile threshold, and where (f) CMPS is greater than its 95-percentile threshold.

From Fig. 8, it is seen that the LVE and O-LVHE models result in nearly identical strain-responses. This can be attributed to the relatively small strain magnitudes (< 5%) under mild rotational acceleration loading, which limits the effect of nonlinearity in the rate-independent part of the stress-strain response (which is linear elastic in the LVE model and Ogden hyperelastic in the O-LVHE model). In the absence of this nonlinearity, LVE and O-LVHE formulations become identical.

Figures 8(c-d) show the time evolution of the brain volume fractions where *E*_*f*_ and MPS are above the 95- percentile MAS and CMPS thresholds (percentile computed from experimental strains), respectively. As expected, the LVE and O-LVHE models predict significantly larger volumes both at the first and second maxima. In terms of relative error, the disagreement between model predictions and experimental observations is more than 100%, which is greater when compared to the relative errors in the peak strain versus time case (Figs. 8(a-b)). For visualization, Figure 8(e) shows the overlap between the predicted and observed brain regions (at *z* = 0 mm slice) where MAS is greater than the MAS95 threshold, for the three investigated constitutive models. From these maps, the O-USS model offers a better accuracy of locating highly strained axonal fibers compared to the LVE and O-LVHE models. From Table 4, the O-USS model leads to an average *D* score for MAS that is ~24% higher compared to that from LVE and O-LVHE models. For the MPS strain, however, there is an insignificant difference between the three models. As shown in Fig. 8(f), while LVE and O-LVHE models predict a larger area above the strain threshold than the experimental observation (i.e., a softer response), O-USS model yields a more conservative estimate; both estimates, however, differ from the experimental response. Similarly, only small differences in the *D* score for the six G-L strain components are observed between the three models.

Lastly, the effect of loading severity on the predicted peak strain maxima of the three models (O-USS, LVE and O-LVHE) is investigated. A sinusoidal rotational acceleration boundary condition is chosen for this purpose (see Fig. 9(a)), which has been commonly employed in the literature to approximate real life loading events and for formulating brain injury criteria^[75,76]^. The associated angular velocity-time curve is also shown in the figure. A fixed time-period of 45 ms is chosen, while the peak angular velocity (*υ*_0_) is increased from 3.42 rad/s to 48 rad/s to simulate an increasing loading severity that can likely cause injury^[75]^.

**Figure 9.**
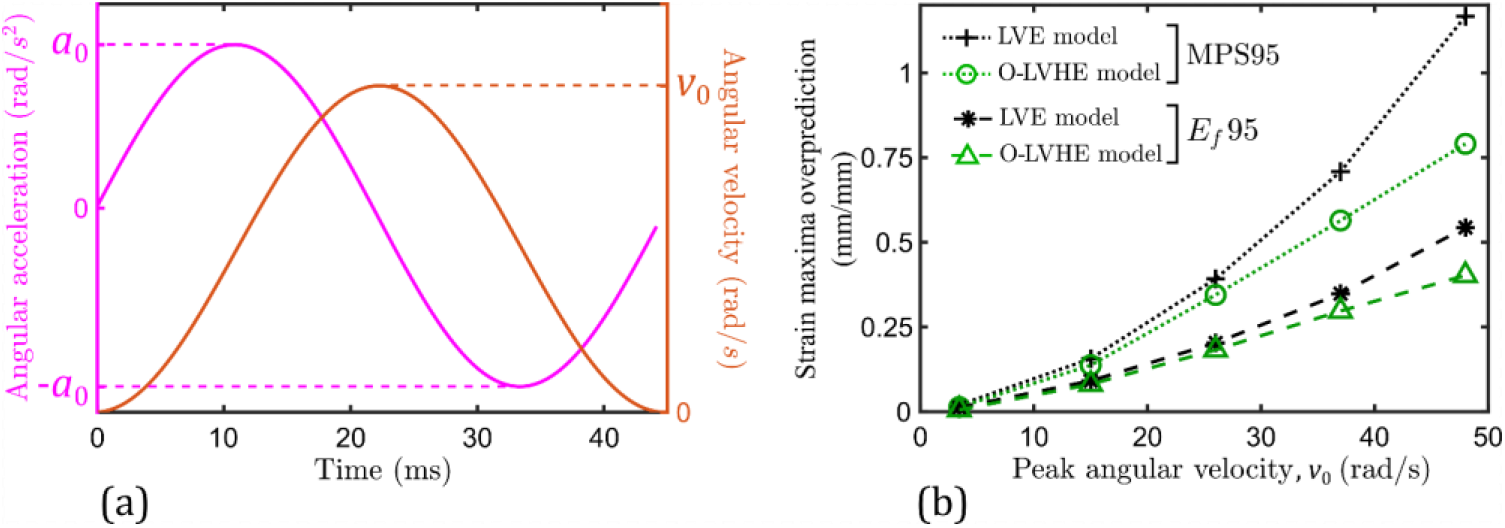
(a) Sinusoidal rotational angular velocity and acceleration versus time boundary condition. (b) Overprediction of the peak (95-percentile) strain maxima of *E*_*f*_ and MPS by the LVE and O-LVHE models (with respect to the O-USS model).

Figure 9(b) plots the overprediction of the maxima of peak *E*_*f*_ and MPS versus time responses by the LVE and O-LVHE models (with respect to the corresponding values from the O-USS model), as a function of applied peak angular velocity. At low peak velocities (3-5 rad/s) such as those employed in the in-vivo tMRI in this study, the predicted magnitude of strain maxima by LVE and O-LVHE models is larger than those predicted by the O-USS model (i.e., a positive strain maxima overprediction). As the loading velocity is increased, the extent of overprediction also increases monotonically, which is attributed to the increasingly nonlinear material response (both rate-independent and rate-sensitive parts) not captured by the LVE and O-LVHE models. Further, unlike low applied velocities that were associated with a nearly identical LVE and O-LVHE strain-response (Fig. 8), at high loading velocities, O-LVHE model predicts smaller strain maxima than the LVE model. This is expected because as strains increase with applied loading velocity, so does the nonlinearity in the rate-independent stresses of the O-LVHE model, resulting in a relatively stiffer brain dynamics than the perfectly linearly elastic case of the LVE model.

## 7. Summary, Discussion, and Limitations

This study proposed a framework for the development and validation of fully-subject specific head models based on a nonlinear visco-hyperelastic constitutive modeling approach. In addition to a personalized head geometry from anatomical images, MRE is employed to derive nonlinear material properties of brain tissue in its native biophysical environment. The head model is used to simulate 3D brain dynamics under mild rotational acceleration to the head about the I/S axis. Full-field strain under the same loading condition are acquired experimentally using tagged MRI, which provides reliable data for model validation. Unlike the previously available head models, the proposed subject-specific modeling framework minimizes uncertainty from the significant variability in head shape and tissue response among the human population. Further, the employment of nonlinear visco-hyperelastic O-USS model allows relating viscous overstress in the material to viscous dissipation at high strain rates, which is unlike the majority of the available head models that employ linear or partially linear visco-hyperelastic constitutive models to represent viscous overstress via a monotonically decreasing relaxation modulus in time. This difference in the modeling philosophy makes this approach especially well-suited for capturing the time-dependent material response at high strain rates^[25]^.

Parameters of the O-USS model are calibrated via a novel in-ex-vivo hybrid parametrization method, which utilizes both in-vivo MRE data and ex-vivo stress-strain data. Ex-vivo data sets are carefully chosen from the literature and incorporated in the optimization process to make sure that (i) the in-vivo tissue mechanics information obtained from MRE is preserved, and at the same time, (ii) the fitted model can capture a wide range of strain and strain rates (or frequencies) that are associated with real-life brain injuries. The obtained model parameters for all the brain substructures feature large absolute values of the compression-tension asymmetry parameter *α* (e.g., 1.41-2.37 kPa for S01) and the nonlinear rate-sensitivity control parameter *k*_21_ (0.35-0.62 kPa.s) and index *c*_21_ (0.58-0.65), providing further evidence of a highly nonlinear mechanical response.

From both the simulation and experiment, it is observed that the various brain substructures undergo peak strains at ~45 ms after head deceleration started. For the dominant, in-plane G-L strain components at this time, a significant overlap is observed between regions under different strain “bins”. Among the two scalar strain measures of MAS and CMPS, a better spatial agreement is observed for regions where MAS is greater than a threshold value; notably, the model accurately located regions of highly strained fibers in the left-frontal and right-posterior regions of the brain. Overall, for the three in-plane G-L strain components and the two scalar strain measures, an average *D* score in the range of 0.40-0.50 is obtained. In addition to good spatial agreement, a good temporal agreement of strains from simulation and experiment is observed, both in terms of the locations of the peak strain as well as the overall dynamics of the strain-time responses. Notably, the simulated peak values of in-plane G-L strain components are within 10% of the corresponding experimental observations; an even better agreement is obtained for *E*_*f*_ and MPS (< 5%). In terms of weighted-average CORA and CS scores, values in the range of 0.42-0.47 and 44-55 are obtained for the in-plane G-L strain components and *E*_*f*_, respectively; for MPS, higher scores of 0.71 and 83.15 are obtained. Across all the validation metrics employed in this study, it is seen that the simulated out-of-plane G-L strains show a relatively poor agreement with experimental observations, which can be attributed to the increased experimental uncertainty and SNR for these small strain values. Although a direct comparison with existing head models from the literature is not possible, considering the currently available “validated” models that show differences with experiments on the order of ±100%^[4,5,12,13,20]^, the head models developed in this work show excellent agreement with experimental data. Note that although results of a single representative subject (S01) are presented, a similarly good agreement between experiment and simulation is seen for two other subjects (see validation scores in the supplementary material), which demonstrates the robustness of the proposed modeling framework in terms of the ability to translate from one subject to another.

The proposed validation approach is also used to quantify the improvement in simulation accuracy provided by the O-USS constitutive model, compared to the commonly used LVE and O-LVHE models. The most notable differences between the three model predictions are seen in the relative errors for the peak strain maxima, and in the overall evolution of the strain-time response; the O-USS model predicts smaller peak strains and a more gradual decay of strain with time compared to the other two models, which agrees well with experimental observations. Note that although the reported comparison in this study employed O-USS model parameters that are “refined” via nonlinear constraint optimization with ex-vivo large deformation experimental stress-strain data, even the head model with initial O-USS model parameter estimates (from step 2 of the in-ex-vivo parametrization) lead to a similar result: smaller peak strains and a gradual strain decay (see supplementary material). This conclusively shows that the improvement in the predicted strain-response is due to the incorporation of nonlinearity in viscous overstress by the O-USS model. The extent of peak strain overprediction by LVE and O-LVHE models compared to the O-USS model increases with the applied rotational velocity or acceleration, pointing to a likely greater disagreement of these two models with the experimental response under potentially injurious loading conditions. The present study recommends the employment of the O-USS constitutive model for the next-generation of computational head models.

Selection of appropriate validation metrics for head injury models has been a matter of debate, with no consensus yet in the brain biomechanics community^[27]^. The present work suggests the employment of three types of validation metrics: *D*, CORA or CS, and relative error, on multiple strain types (**E**, *E*_*f*_ and MPS). Notably, these metrics are complementary and not necessarily correlated, i.e., they analyze different aspects of the strain-response, and a good validation score in one category (say, spatial) does not guarantee a good score in the other category (say, temporal). This quality allowed clear discrimination of the O-USS-, LVE- and O-LVHE-based models; here, *D* scores showed similar spatial accuracy for all the investigated strain measures except MAS, whereas the CORA and CS scores resulted in consistently greater temporal scores for the O-USS-based model, suggesting improvement in predicted strain-time evolution. At the same time, incorporation of tensorial strain in addition to the commonly used scalar strain measures can prevent models with very different strain tensor predictions (which can lead to same scalar *E*_*f*_ and MPS), from leading to similar validation scores^[27]^.

Due to the nature of MRE that relies on estimating properties from imaged shear wave propagation, it is not possible to estimate the properties of small and very stiff structures such as falx, tentorium, and SAS. In the present study, material properties of these substructures are derived from ex-vivo studies, and thus are not subject-specific. Furthermore, MRE only probes the material response in a limited frequency range. The in-ex-vivo hybrid parametrization procedure developed in this work overcomes this limitation to an extent by including ex-vivo experimental data; still, it is anticipated that broadening the harmonic excitation frequency range is a possibility, which will be explored in the future work.

Head injury models in the literature have almost exclusively been validated using high rate cadaveric impact experiments^[7,8]^, which are associated with a very short (3–5 ms) impulse duration and a peak rotational acceleration of over 7.9 krad/s^2^. Unlike these models, the present subject-specific head model is validated at non-injurious mild rotational accelerations, which have a longer loading duration of ~40–50 ms, and a peak rotational acceleration of ~150–350 rad/s^2^. A recent study^[27]^ has noted the importance of both of these extremes of loading severity in the validation of computational head injury models. This is because head kinematics during many of the common real-life head injuries fall in-between these two extremes. However, due to human subject safety considerations, a subject-specific validation at injurious rotational accelerations is not possible. The present study is of the opinion that high resolution, time-varying, and full-field strain data from cadaveric experiments (currently unavailable) may still be valuable for validating subject-specific head models using the validation approach of this study, even though with a lower confidence because of the potential differences in material properties between live and cadaveric brain tissue. This will be the subject of future research.

## Acknowledgements

This research was supported by the National Institute of Neurological Disorders and Stroke of the National Institutes of Health under Award Number U01NS112120. The content is solely the responsibility of the authors and does not necessarily represent the official views of the National Institutes of Health.

## Appendix A: Viscous Dissipation Potential Model Parameter Refinement

The following nonlinear constrained optimization problem is employed to obtain “refined” viscous dissipation model parameters:

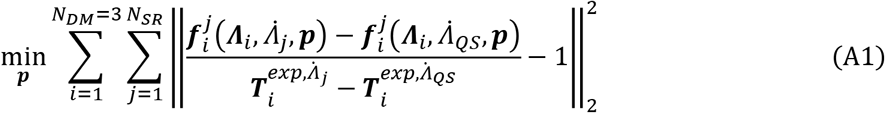

where ***p*** = [*k*_11_, *k*_21_, *c*_21_]^T^ is the vector of O-USS model parameters to be optimized, *i* represents a particular deformation mode (DM) (*i* = 1, 2 and 3 are shear, compression and tension modes, respectively), and *j* represents a particular dynamic strain rate level, *N*_*SR*_ being the number of applied dynamic strain rates in the ex-vivo study (not including the quasi-static (QS) strain rate). 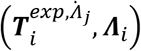 is the experimentally obtained nominal stress and strain (*∧* can be shear strain *γ* (for *i* = 1) or uniaxial strain *ε* (for *i* = 2 and 3)) dataset for the *i*^*th*^ deformation mode and at the *j*^*th*^ dynamic strain rate level; 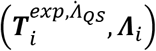 is the corresponding dataset at the lowest strain rate level, 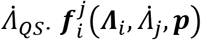 is the vector containing symbolic nominal stress as formulated using the O-USS model (Eqs. (6–9)); note, nominal stress is *J***F**^−1^**σ** (where **σ** is the Cauchy stress). 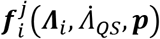 are the corresponding nominal stress equations for the lowest strain rate level, 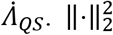 represents the square of the 2-norm. The following constraints are put on the optimization problem solution space:

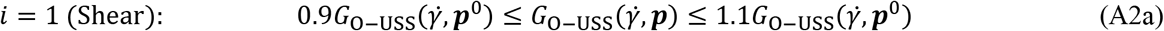

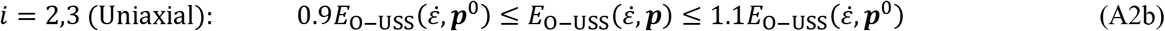

where 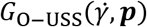 and 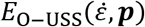 are the symbolic expressions of tangent shear and uniaxial moduli of the O-USS model, respectively, evaluated for the parameter set ***p*** (given in Eq. (14)) and ***p***^0^ is the set of model parameters, 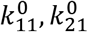, and 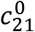, obtained in the previous step. 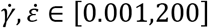 (same range is considered for computation of the apparent elastic moduli-strain rate spectra). The optimization problem in Eqs. (A1–A2) are solved in MATLAB for each brain voxel using the *fmincon* function (via interior-point algorithm) with 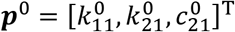 as the initial guess.

## References

[1] “National Center for Health Statistics: Mortality Data on CDC WONDER,” can be found under https://wonder.cdc.gov/mcd.html, **n.d.**

[2] S. Ji, in Encycl. Comput. Neurosci., Springer New York, New York, NY, 2018, pp. 1–4.

[3] A. Madhukar, M. Ostoja-Starzewski, Ann. Biomed. Eng. 2019, 47, 1832.

[4] D. Sahoo, C. Deck, R. Willinger, J. Mech. Behav. Biomed. Mater. 2014, 33, 24.

[5] S. Ji, W. Zhao, J. C. Ford, J. G. Beckwith, R. P. Bolander, R. M. Greenwald, L. A. Flashman, K. D. Paulsen, T. W. McAllister, J. Neurotrauma 2015, 32, 441.

[6] S. Ganpule, N. P. Daphalapurkar, K. T. Ramesh, A. K. Knutsen, D. L. Pham, P. V. Bayly, J. L. Prince, J. Neurotrauma 2017, 34, 2154.

[7] W. N. Hardy, C. D. Foster, M. J. Mason, K. H. Yang, A. I. King, S. Tashman, Stapp Car Crash J. 2001, 45, 337.

[8] W. N. Hardy, M. J. Mason, C. D. Foster, C. S. Shah, J. M. Kopacz, K. H. Yang, A. I. King, J. Bishop, M. Bey, W. Anderst, S. Tashman, Stapp Car Crash J. 2007, 51, 17.

[9] D. B. MacManus, B. Pierrat, J. G. Murphy, M. D. Gilchrist, Sci. Rep. 2017, 7, 13729.

[10] M. T. Prange, D. F. Meaney, S. S. Margulies, Stapp Car Crash J. 2000, 44, 205.

[11] S. Chatelin, A. Constantinesco, R. Willinger, Biorheology 2010, 47, 255.

[12] C. Giordano, S. Kleiven, Stapp Car Crash J. 2014, 58, 29.

[13] H. Mao, L. Zhang, B. Jiang, V. V. Genthikatti, X. Jin, F. Zhu, R. Makwana, A. Gill, G. Jandir, A. Singh, K. H. Yang, J. Biomech. Eng. 2013, 135, DOI 10.1115/1.4025101.

[14] S. Budday, T. C. Ovaert, G. A. Holzapfel, P. Steinmann, E. Kuhl, Arch. Comput. Methods Eng. 2020, 27, 1187.

[15] L. A. Mihai, S. Budday, G. A. Holzapfel, E. Kuhl, A. Goriely, J. Mech. Phys. Solids 2017, 106, 60.

[16] L. E. Miller, J. E. Urban, J. D. Stitzel, Biomech. Model. Mechanobiol. 2016, 15, 1201.

[17] B. Yang, K. M. Tse, N. Chen, L. Bin Tan, Q. Q. Zheng, H. M. Yang, M. Hu, G. Pan, H. P. Lee, Biomed Res. Int. 2014, 2014, DOI 10.1155/2014/408278.

[18] S. Budday, G. Sommer, C. Birkl, C. Langkammer, J. Haybaeck, J. Kohnert, M. Bauer, F. Paulsen, P. Steinmann, E. Kuhl, G. A. Holzapfel, Acta Biomater. 2017, 48, 319.

[19] K. Upadhyay, G. Subhash, D. Spearot, J. Mech. Phys. Solids 2020, 135, 103777.

[20] T. Wu, A. Alshareef, J. S. Giudice, M. B. Panzer, Ann. Biomed. Eng. 2019, 47, 1908.

[21] S. Kleiven, Stapp Car Crash J. 2007, 51, 81.

[22] S. Nicolle, M. Lounis, R. Willinger, J.-F. Palierne, Biorheology 2005, 42, 209.

[23] M. Hrapko, J. A. W. van Dommelen, G. W. M. Peters, J. S. H. M. Wismans, Biorheology 2006, 43, 623.

[24] G. Peters, J. Meulman, A. Sauren, Biorheology 1997, 34, 127.

[25] D. P. Pioletti, in Mech. Biol. Tissue, Springer-Verlag, Berlin/Heidelberg, 1999, pp. 399–404.

[26] R. W. Ogden, Proc. R. Soc. London. A. Math. Phys. Sci. 1972, 326, 565.

[27] W. Zhao, B. Choate, S. Ji, J. Mech. Behav. Biomed. Mater. 2018, 80, 222.

[28] P. V. Bayly, A. Alshareef, A. K. Knutsen, K. Upadhyay, R. J. Okamoto, A. Carass, J. A. Butman, D. L. Pham, J. L. Prince, K. T. Ramesh, C. L. Johnson, Ann. Biomed. Eng. 2021, DOI 10.1007/s10439-021-02820-0.

[29] A. Alshareef, A. K. Knutsen, C. L. Johnson, A. Carass, K. Upadhyay, P. V Bayly, D. L. Pham, J. L. Prince, K. T. Ramesh, Brain Multiphysics 2021, 100038.

[30] Y. Huo, A. J. Plassard, A. Carass, S. M. Resnick, D. L. Pham, J. L. Prince, B. A. Landman, Neuroimage 2016, 138, 197.

[31] J. Glaister, M. Shao, X. Li, A. Carass, S. Roy, A. M. Blitz, J. L. Prince, L. Ellingsen, Proc SPIE 2018, 10574, 1057431.

[32] J. Glaister, A. Carass, D. L. Pham, J. A. Butman, J. L. Prince, Med Image Comput Comput Assist Interv 2017, 10433, 92.

[33] P. A. Yushkevich, J. Piven, H. C. Hazlett, R. G. Smith, S. Ho, J. C. Gee, G. Gerig, Neuroimage 2006, 31, 1116.

[34] A. Shiino, Y. Chen, K. Tanigaki, A. Yamada, P. Vigers, T. Watanabe, I. Tooyama, I. Akiguchi, Sci. Rep. 2017, 7, 39818.

[35] B. B. Avants, N. J. Tustison, M. Stauffer, G. Song, B. Wu, J. C. Gee, Front. Neuroinform. 2014, 8, 1.

[36] S. Roy, J. A. Butman, D. L. Pham, Neuroimage 2017, 146, 132.

[37] L. V Hiscox, H. Schwarb, M. D. J. McGarry, C. L. Johnson, Neuroimage 2021, 232, 117889.

[38] M. D. J. McGarry, E. E. W. Van Houten, C. L. Johnson, J. G. Georgiadis, B. P. Sutton, J. B. Weaver, K. D. Paulsen, Med. Phys. 2012, 39, 6388.

[39] L. V. Hiscox, C. L. Johnson, E. Barnhill, M. D. J. McGarry, J. Huston, E. J. R. van Beek, J. M. Starr, N. Roberts, Phys. Med. Biol. 2016, 61, R401.

[40] A. D. Gomez, A. K. Knutsen, F. Xing, Y.-C. Lu, D. Chan, D. L. Pham, P. Bayly, J. L. Prince, IEEE Trans. Biomed. Eng. 2019, 66, 1456.

[41] A. K. Knutsen, A. D. Gomez, M. Gangolli, W. Wang, D. Chan, Y. Lu, E. Christoforou, J. L. Prince, P. V Bayly, J. A. Butman, D. L. Pham, Brain Multiphysics 2020, 100015.

[42] C. Pierpaoli, L. Walker, M. O. Irfanoglu, A. Barnett, P. Basser, L.-C. Chang, C. Koay, S. Pajevic, G. Rohde, J. Sarlls, M. Wu, in Proc. Int. Soc. Magn. Reson. Med., Stockholm, Sweden, 2010.

[43] S. Wakana, L. M. Nagae-Poetscher, H. Jiang, P. van Zijl, X. Golay, S. Mori, Magn. Reson. Med. 2005, 53, 649.

[44] D. P. Pioletti, L. R. Rakotomanana, J. F. Benvenuti, P. F. Leyvraz, J. Biomech. 1998, 31, 753.

[45] A. I. Zhurov, G. Limbert, D. P. Aeschlimann, J. Middleton, Comput. Methods Biomech. Biomed. Engin. 2007, 10, 223.

[46] J. C. Simo, C. Miehe, Comput. Methods Appl. Mech. Eng. 1992, 98, 41.

[47] S. Ganpule, N. P. Daphalapurkar, M. P. Cetingul, K. T. Ramesh, Shock Waves 2018, 28, 127.

[48] R. J. H. Cloots, J. A. W. van Dommelen, T. Nyberg, S. Kleiven, M. G. D. Geers, Biomech. Model. Mechanobiol. 2011, 10, 413.

[49] L. A. Mihai, L. Chin, P. A. Janmey, A. Goriely, J. R. Soc. Interface 2015, 12, 20150486.

[50] K. Upadhyay, G. Subhash, D. Spearot, Int. J. Eng. Sci. 2020, 154, 103314.

[51] K. Upadhyay, G. Subhash, D. Spearot, J. Mech. Phys. Solids 2019, 124, 115.

[52] K. Upadhyay, D. Spearot, G. Subhash, Int. J. Impact Eng. 2021, 156, 103949.

[53] X. Jin, F. Zhu, H. Mao, M. Shen, K. H. Yang, J. Biomech. 2013, 46, 2795.

[54] J. R. Macdonald, Rev. Mod. Phys. 1966, 38, 669.

[55] M. Bulat, in Intracranial Press. VIII, Springer Berlin Heidelberg, Berlin, Heidelberg, 1993, pp. 726–730.

[56] R. H. Cole, Underwater Explosions., Princeton Univ. Press, Princeton, 1948.

[57] S. Kleiven, Stapp Car Crash J. 2007, 51, 81.

[58] G. Mattei, A. Ahluwalia, Mech. Time-Dependent Mater. 2019, 23, 327.

[59] L. Bartolini, D. Iannuzzi, G. Mattei, Sci. Rep. 2018, 8, 1.

[60] A. Tirella, G. Mattei, A. Ahluwalia, J. Biomed. Mater. Res. - Part A 2014, 102, 3352.

[61] D. Sahoo, C. Deck, R. Willinger, Accid. Anal. Prev. 2016, 92, 53.

[62] Y.-C. Lu, N. P. Daphalapurkar, A. K. Knutsen, J. Glaister, D. L. Pham, J. A. Butman, J. L. Prince, P. V. Bayly, K. T. Ramesh, Ann. Biomed. Eng. 2019, 47, 1923.

[63] J. E. Galford, J. H. McElhaney, J. Biomech. 1970, 3, 211.

[64] J. H. McElhaney, J. W. Melvin, V. L. Roberts, H. D. Portnoy, in Perspect. Biomed. Eng. (Ed.: R.M. Kenedi), Palgrave Macmillan UK, London, 1973, pp. 215–222.

[65] W. Goldsmith, in Biomech. Its Found. Object. (Ed.: Y. Fung), Prentice Hall, Englewood Cliffs, NJ, 1972, pp. 585–634.

[66] D. Sulsky, Z. Chen, H. L. Schreyer, Comput. Methods Appl. Mech. Eng. 1994, 118, 179.

[67] D. Sulsky, S. J. Zhou, H. L. Schreyer, Comput. Phys. Commun. 1995, 87, 236.

[68] J. L. González Acosta, P. J. Vardon, G. Remmerswaal, M. A. Hicks, Comput. Mech. 2020, 65, 555.

[69] N. S. Pruijn, The Improvement of the Material Point Method by Increasing Efficiency and Accuracy, Delft University ofTechnology, 2016.

[70] W. Zhao, Y. Cai, Z. Li, S. Ji, Biomech. Model. Mechanobiol. 2017, 16, 1709.

[71] C. Gehre, H. Gades, P. Wernicke, in Proc. 21ST Int. Tech. Conf. Enhanc. Saf. Veh., National Highway Traffic Safety Administration, Stuttgart, Germany, 2009, pp. 1–8.

[72] H. Kimpara, Y. Nakahira, M. Iwamoto, K. Miki, K. Ichihara, S. ichi Kawano, T. Taguchi, Stapp Car Crash J. 2006, 50, 509.

[73] C. Giordano, S. Kleiven, Stapp Car Crash J. 2016, 60, 363.

[74] O. Starkova, A. Aniskevich, Mech. Time-Dependent Mater. 2007, 11, 111.

[75] E. G. Takhounts, M. J. Craig, K. Moorhouse, J. McFadden, V. Hasija, in Stapp Car Crash J., 2013, pp. 243–266.

[76] L. F. Gabler, J. R. Crandall, M. B. Panzer, Ann. Biomed. Eng. 2018, 46, 972.

