## Supplemental Material for "Development and Validation of Subject-Specific 3D Human Head Models Based on a Nonlinear Visco-Hyperelastic Constitutive Framework"

### Supplementary material

### SM1. Derivation of the O-USS model

From Section 2.2 of the main manuscript, the general form of a viscous dissipation-based visco-hyperelastic constitutive model is

$$\boldsymbol{\sigma} = \boldsymbol{\sigma}_{\text{vol}} + \boldsymbol{\sigma}_{\text{iso}} = \boldsymbol{\sigma}_{\text{vol}} + \boldsymbol{\sigma}_{h,\text{iso}} + \boldsymbol{\sigma}_{v,\text{iso}} \tag{1}$$

with

$$\boldsymbol{\sigma}_{\text{vol}} = \frac{2}{J}\mathbf{F}\frac{\partial U(J)}{\partial \mathbf{C}}\mathbf{F}^{\text{T}}, \qquad \boldsymbol{\sigma}_{h,\text{iso}} = \frac{2}{J}\mathbf{F}\frac{\partial \bar{W}_h(\bar{\mathbf{C}}, \mathbf{A}_0)}{\partial \mathbf{C}}\mathbf{F}^{\text{T}}, \qquad \boldsymbol{\sigma}_{v,\text{iso}} = \frac{2}{J}\mathbf{F}\frac{\partial \bar{W}_v\left(\bar{\mathbf{C}}, \dot{\bar{\mathbf{C}}}, \mathbf{A}_0\right)}{\partial \dot{\mathbf{C}}}\mathbf{F}^{\text{T}} \tag{2}$$

In Eq. (1), Cauchy stress $\boldsymbol{\sigma}$ is additively decomposed into separate volumetric ($\boldsymbol{\sigma}_{\text{vol}}$) and isochoric ($\boldsymbol{\sigma}_{\text{iso}}$) components. The latter is further split into hyperelastic ($\boldsymbol{\sigma}_{h,\text{iso}}$) and viscous overstress ($\boldsymbol{\sigma}_{v,\text{iso}}$) components. Specifically, the rate-independent volumetric stress is derived from the volumetric energy density $U$, which is a function of $J = \det(\mathbf{F})$ ($\mathbf{F}$ is the deformation gradient tensor). $\mathbf{C} = \mathbf{F}^{\text{T}}\mathbf{F}$ is the right Cauchy-Green deformation tensor. Further, the isochoric hyperelastic stress $\boldsymbol{\sigma}_{h,\text{iso}}$ that is associated with the quasi-static material response is derived from a strain energy density function $\bar{W}_h$, which depends on the modified right Cauchy-Green deformation tensor $\bar{\mathbf{C}}$ ($\bar{\mathbf{C}} = J^{-2/3}\mathbf{C}$) and the structure tensor $\mathbf{A}_0$. ). Finally, the viscous overstress $\boldsymbol{\sigma}_{v,\text{iso}}$ is associated with the dynamic rate-dependent material response and is derived using a viscous dissipation potential $\bar{W}_v$, which is a function of $\bar{\mathbf{C}}, \dot{\bar{\mathbf{C}}}$ and $\mathbf{A}_0$.

In the present study, the Siemo-Miehe form of volumetric energy density, the Ogden hyperelastic strain energy density, and the Upadhyay-Subhash-Spearot (USS) viscous dissipation potential are considered (all these forms assume isotropic material response):

$$U(J)=\frac{\kappa}{2}\left(\frac{J^2-1}{2}-\ln J\right) \tag{3}$$

$$\bar{W}_h(\bar{\mathbf{C}})=\bar{W}_h(\bar{\lambda}_1,\bar{\lambda}_2,\bar{\lambda}_3)=\frac{2\mu_\infty}{\alpha^2}\left(\bar{\lambda}_1^\alpha+\bar{\lambda}_2^\alpha+\bar{\lambda}_3^\alpha-3\right) \tag{4}$$

$$\bar{W}_v\left(\bar{\mathbf{C}},\dot{\bar{\mathbf{C}}}\right)=\bar{W}_v(\bar{I}_1,\bar{I}_2,\bar{J}_2,\bar{J}_5)=k_{11}\bar{J}_2\sqrt{\bar{I}_1-3}+\frac{k_{21}}{c_{21}}\bar{J}_5^{c_{21}}\sqrt{\bar{I}_2-3} \tag{5}$$

where $\kappa$, $\mu_\infty$, $\alpha$, $k_{11}$, $k_{21}$ and $c_{21}$ are model parameters. Further, $\bar{\lambda}_1$, $\bar{\lambda}_2$ and $\bar{\lambda}_3$ are the distortional principal stretches, such that $\bar{\mathbf{C}}=\sum_{i=1}^{3}\bar{\lambda}_i^2\widehat{\boldsymbol{N}}_i\otimes\widehat{\boldsymbol{N}}_i$ (here, $\widehat{\boldsymbol{N}}_i$ are the eigen vectors of $\bar{\mathbf{C}}$, also called principal referential directions[1]). Note, $\widehat{\boldsymbol{N}}_i$ are also the eigen vectors of the second Piola-Kirchhoff stress tensor. Equivalently, the modified left Cauchy-Green deformation tensor can be written in its spectral form as $\bar{\mathbf{B}}=\bar{\mathbf{F}}\bar{\mathbf{F}}^{\mathrm{T}}=\sum_{i=1}^{3}\bar{\lambda}_i^2\widehat{\boldsymbol{n}}_i\otimes\widehat{\boldsymbol{n}}_i$ (here, $\widehat{\boldsymbol{n}}_i$ are the eigen vectors of $\bar{\mathbf{B}}$, also called principal spatial directions[1]). $\widehat{\boldsymbol{n}}_i$ are also the eigen vectors of the Cauchy stress, $\boldsymbol{\sigma}$. Notice that $\bar{\mathbf{C}}$ and $\bar{\mathbf{B}}$ have the same eigen values but different eigen vectors. Finally, $\bar{I}_1=\mathrm{tr}(\bar{\mathbf{C}})$ and $\bar{I}_2=1/2\left(\bar{I}_1^2-\mathrm{tr}(\bar{\mathbf{C}}^2)\right)$ are principal invariants of $\bar{\mathbf{C}}$, and $\bar{J}_2=\mathrm{tr}\left(\dot{\bar{\mathbf{C}}}^2\right)$ and $\bar{J}_5=\mathrm{tr}\left(\bar{\mathbf{C}}\dot{\bar{\mathbf{C}}}^2\right)$ are the scalar invariants of tensors $\bar{\mathbf{C}}$ and $\dot{\bar{\mathbf{C}}}$.

For the volumetric energy density in Eq. (3), the volumetric stress component in Eq. (2) can be computed using the chain rule of differentiation as

$$\boldsymbol{\sigma}_{\mathrm{vol}}=\frac{2}{J}\mathbf{F}\left[\frac{dU(J)}{dJ}\frac{\partial J}{\partial\mathbf{C}}\right]\mathbf{F}^{\mathrm{T}} \tag{6}$$

where

$$\frac{dU(J)}{dJ}=\frac{\kappa}{2}\left(J-\frac{1}{J}\right) \tag{7a}$$

$$\frac{\partial J}{\partial\mathbf{C}}=\frac{\partial(\det(\mathbf{C}))^{\frac{1}{2}}}{\partial\mathbf{C}}=\frac{1}{2}J\mathbf{C}^{-1} \tag{7b}$$

From Eqs. (6) and (7), the volumetric stress component is obtained as

$$\boldsymbol{\sigma}_{\mathrm{vol}}=\frac{\kappa}{2}\left(J-\frac{1}{J}\right)\mathbf{I} \tag{8}$$

where $\mathbf{I}$ is the unit symmetric tensor.

Among the Ogden strain energy density and USS viscous dissipation potential, only the latter is based on invariants forming the integrity basis of the tensors $\bar{\mathbf{C}}$ and $\dot{\bar{\mathbf{C}}}$. The Ogden model, on the other hand, is based on the principal stretches of $\bar{\mathbf{C}}$. For such models ($\bar{W}_h(\bar{\mathbf{C}})=\bar{W}_h(\bar{\lambda}_1,\bar{\lambda}_2,\bar{\lambda}_3)$), the isochoric hyperelastic stress component in Eq. (2) can be written as[1]

$$\boldsymbol{\sigma}_{h,\mathrm{iso}}=\frac{2}{J}\mathbf{F}\frac{\partial\bar{W}_h(\bar{\mathbf{C}})}{\partial\mathbf{C}}\mathbf{F}^{\mathrm{T}}=\sum_{i=1}^{3}J^{-1}\left(\bar{\lambda}_i\frac{\partial\bar{W}_h(\bar{\lambda}_i)}{\partial\bar{\lambda}_i}\right)\widehat{\boldsymbol{n}}_i\otimes\widehat{\boldsymbol{n}}_i \tag{9}$$

After computing the derivative of the Ogden strain energy density in Eq. (4) with respect to $\bar{\lambda}_i$, and substituting that expression in Eq. (9), the isochoric hyperelastic stress component is obtained as

$$\boldsymbol{\sigma}_{h,\text{iso}} = \sum_{i=1}^{3} J^{-1}\left(\frac{2\mu_\infty}{\alpha}\left(\bar{\lambda}_i^\alpha - \frac{1}{3}\left[\bar{\lambda}_1^\alpha + \bar{\lambda}_2^\alpha + \bar{\lambda}_3^\alpha\right]\right)\right)\hat{\boldsymbol{n}}_i \otimes \hat{\boldsymbol{n}}_i \tag{10}$$

Finally, for the USS viscous dissipation potential in Eq. (5), the isochoric viscous overstress component in Eq. (2) can be computed using the chain rule of differentiation as

$$\boldsymbol{\sigma}_{v,\text{iso}} = \frac{2}{J}\mathbf{F}\left[\sum_{m=2,5}\frac{\partial \bar{W}_v(\bar{I}_1,\bar{I}_2,\bar{J}_2,\bar{J}_5)}{\partial \bar{J}_m}\left(\left(\frac{\partial \dot{\bar{\mathbf{C}}}}{\partial \dot{\mathbf{C}}}\right)^T : \frac{\partial \bar{J}_m}{\partial \dot{\bar{\mathbf{C}}}}\right)\right]\mathbf{F}^{\mathrm{T}} \tag{11}$$

where

$$\frac{\partial \bar{W}_v(\bar{I}_1,\bar{I}_2,\bar{J}_2,\bar{J}_5)}{\partial \bar{J}_2} = k_{11}\sqrt{\bar{I}_1 - 3};\ \frac{\partial \bar{W}_v(\bar{I}_1,\bar{I}_2,\bar{J}_2,\bar{J}_5)}{\partial \bar{J}_5} = k_{21}\bar{J}_5^{c_{21}-1}\sqrt{\bar{I}_2 - 3} \tag{12$a$}$$

$$\frac{\partial \dot{\bar{\mathbf{C}}}}{\partial \dot{\mathbf{C}}} = \frac{\partial \bar{\mathbf{C}}}{\partial \mathbf{C}} = \frac{\partial J^{-2/3}\mathbf{C}}{\partial \mathbf{C}} = J^{-2/3}\mathbb{I} + \mathbf{C}\otimes\frac{\partial J^{-2/3}}{\partial \mathbf{C}} = J^{-2/3}\left(\mathbb{I} - \frac{1}{3}\mathbf{C}\otimes\mathbf{C}^{-1}\right) = J^{-2/3}\mathbb{P}^{\mathrm{T}} \tag{12$b$}$$

$$\frac{\partial \bar{J}_2}{\partial \dot{\bar{\mathbf{C}}}} = 2\dot{\bar{\mathbf{C}}};\ \frac{\partial \bar{J}_5}{\partial \dot{\bar{\mathbf{C}}}} = \bar{\mathbf{C}}\dot{\bar{\mathbf{C}}} + \dot{\bar{\mathbf{C}}}\bar{\mathbf{C}} \tag{12$c$}$$

Note, for the proof of equivalence of $\frac{\partial \dot{\bar{\mathbf{C}}}}{\partial \dot{\mathbf{C}}}$ and $\frac{\partial \bar{\mathbf{C}}}{\partial \mathbf{C}}$ in Eq. (12b), readers are referred to Zhurov et al.[2]. $\mathbb{P}$ is called the "referential fourth order projection tensor",

$$\mathbb{P} \coloneqq \mathbb{I} - \frac{1}{3}\mathbf{C}^{-1}\otimes\mathbf{C} \tag{13}$$

where $\mathbb{I}$ is the fourth order unit tensor (in index notation, $\mathbb{I}_{ijkl} = \delta_{ik}\delta_{jl}$, where $\delta$ is the Kronecker delta symbol). After substituting Eqs. 12(a-c) into Eq. (11) and simplifying, the isochoric viscous overstress component is obtained as

$$\boldsymbol{\sigma}_{v,\text{iso}} = J^{-1}\left(8k_{11}\sqrt{\bar{I}_1 - 3}\,\text{dev}(\bar{\mathbf{B}}\bar{\mathbf{D}}\bar{\mathbf{B}}) + 4k_{21}\bar{J}_5^{c_{21}-1}\sqrt{\bar{I}_2 - 3}\,\text{dev}(\bar{\mathbf{B}}\cdot(\bar{\mathbf{B}}\bar{\mathbf{D}}\bar{\mathbf{B}}) + (\bar{\mathbf{B}}\bar{\mathbf{D}}\bar{\mathbf{B}})\cdot\bar{\mathbf{B}})\right) \tag{14}$$

where $\text{dev}(\cdot) = (\cdot) - \frac{1}{3}\text{tr}(\cdot)\mathbf{I}$ is the deviatoric operator in the Eulerian description, and $\bar{\mathbf{D}}$ is the modified rate of deformation tensor, which is the symmetric part of the modified spatial velocity gradient $\bar{\mathbf{L}}$ (i.e., $\bar{\mathbf{D}} = (\bar{\mathbf{L}} + \bar{\mathbf{L}}^{\mathrm{T}})/2$; where $\bar{\mathbf{L}} = \dot{\bar{\mathbf{F}}}\bar{\mathbf{F}}^{\mathrm{T}}$). For more information on the characterization of Cauchy stress tensor in terms of the projection tensor, see Holzapfel[1].

Using Eqs. (8), (10) and (14), the total stress in the O-USS constitutive model is expressed as

$$\boldsymbol{\sigma} = \frac{\kappa}{2}\left(J - \frac{1}{J}\right)\mathbf{I} + \sum_{i=1}^{3} J^{-1}\left(\frac{2\mu_\infty}{\alpha}\left(\bar{\lambda}_i^\alpha - \frac{1}{3}\left[\bar{\lambda}_1^\alpha + \bar{\lambda}_2^\alpha + \bar{\lambda}_3^\alpha\right]\right)\right)\hat{\boldsymbol{n}}_i \otimes \hat{\boldsymbol{n}}_i +$$
$$J^{-1}\left(8k_{11}\sqrt{\bar{I}_1 - 3}\,\text{dev}(\bar{\mathbf{B}}\bar{\mathbf{D}}\bar{\mathbf{B}}) + 4k_{21}\bar{J}_5^{c_{21}-1}\sqrt{\bar{I}_2 - 3}\,\text{dev}(\bar{\mathbf{B}}\cdot(\bar{\mathbf{B}}\bar{\mathbf{D}}\bar{\mathbf{B}}) + (\bar{\mathbf{B}}\bar{\mathbf{D}}\bar{\mathbf{B}})\cdot\bar{\mathbf{B}})\right) \tag{6}$$

### SM2. LVE model parameters of different brain substructures for subjects S01, S02 and S03

**Table 1**. Average linear viscoelastic model parameters of the different brain substructures for subject S01 (31/M).

| Brain substructure | LVE model parameters | | | | |
|---|---|---|---|---|---|
| | $G_0$ (kPa) | $g_1$ | $\tau_1$ (ms) | $g_2$ | $\tau_2$ (ms) |
| Deep gray matter | 8.22 | 0.75 | 0.94 | 0.05 | 29.30 |
| Cortical gray matter | 5.72 | 0.74 | 1.00 | 0.05 | 29.60 |
| Corpus Callosum | 7.04 | 0.68 | 1.19 | 0.05 | 30.82 |
| Corona Radiata | 8.13 | 0.76 | 0.90 | 0.04 | 28.90 |
| Cerebellum gray matter | 4.44 | 0.63 | 1.39 | 0.06 | 31.59 |
| Cerebellum white matter | 6.03 | 0.64 | 1.37 | 0.05 | 31.54 |
| Brainstem | 5.44 | 0.62 | 1.23 | 0.07 | 24.73 |

**Table 2**. Average linear viscoelastic model parameters of the different brain substructures for subject S02 (46/M).

| Brain substructure | LVE model parameters | | | | | | |
|---|---|---|---|---|---|---|---|
| | $G_0$ (kPa) | $g_1$ | $\tau_1$ (ms) | $g_2$ | $\tau_2$ (ms) | $g_3$ | $\tau_3$ (ms) |
| Deep gray matter | 8.83 | 0.01 | 62.55 | 0.26 | 0.78 | 0.48 | 0.57 |
| Cortical gray matter | 9.04 | 0.01 | 64.27 | 0.39 | 0.64 | 0.40 | 0.62 |
| Corpus Callosum | 6.74 | 0 | 20.65 | 0.40 | 1.16 | 0.30 | 0.15 |
| Corona Radiata | 12.61 | 0.01 | 69.75 | 0.81 | 0.72 | 0.01 | 0.70 |
| Cerebellum gray matter | 5.41 | 0.01 | 48.02 | 0.43 | 1.06 | 0.26 | 0.62 |
| Cerebellum white matter | 7.97 | 0 | 44.70 | 0.41 | 0.96 | 0.32 | 0.95 |
| Brainstem | 8.45 | 0.02 | 10.98 | 0.23 | 1.04 | 0.46 | 1.03 |

**Table 3**. Average linear viscoelastic model parameters of the different brain substructures for subject S03 (28/M).

| Brain substructure | LVE model parameters | | | | | | |
|---|---|---|---|---|---|---|---|
| | $G_0$ (kPa) | $g_1$ | $\tau_1$ (ms) | $g_2$ | $\tau_2$ (ms) | $g_3$ | $\tau_3$ (ms) |
| Deep gray matter | 8.07 | 0.02 | 50.63 | 0.64 | 0.97 | 0.07 | 0.93 |
| Cortical gray matter | 7.50 | 0.03 | 44.68 | 0.42 | 0.87 | 0.33 | 0.64 |
| Corpus Callosum | 8.92 | 0.02 | 47.09 | 0.72 | 0.86 | 0.03 | 0.60 |
| Corona Radiata | 8.48 | 0.03 | 42.19 | 0.39 | 0.73 | 0.36 | 0.70 |
| Cerebellum gray matter | 4.92 | 0.05 | 48.55 | 0.33 | 1.38 | 0.29 | 0.76 |
| Cerebellum white matter | 6.30 | 0.05 | 45.24 | 0.41 | 1.07 | 0.27 | 0.74 |
| Brainstem | 5.26 | 0.05 | 48.69 | 0.65 | 0.99 | 0.03 | 1.58 |

### SM3. Ex-vivo studies selected for the hybrid in-ex-vivo parameterization

**Table 4**. Summary of the ex-vivo studies selected for the hybrid in-ex-vivo parameterization of each brain substructure in this study.

| Brain substructure | Ex-vivo study | | | | |
|---|---|---|---|---|---|
| | Reference | Applied maximum strain (mm/mm) | Deformation modes | Applied strain rates ($s^{-1}$) | Subject species |
| Deep gray matter; Cortical gray matter; Corpus Callosum; Corona Radiata | Jin et al. (2013)[33] | 0.5 | Compression, tension, shear | 0.5, 5, 30 | Human |
| Cerebellum gray matter; Cerebellum white matter; Brainstem | Li et al. (2020)[75] | 0.5 | Compression, tension, shear | 0.01, 1, 50 | Pig (pediatric) |

### SM4. O-USS model parameters for subjects S02 and S03

**Table 5**. O-USS material properties of the various brain substructures for subject S02 (46/M).

| Brain substructure | Material properties (O-USS model) | | | | | | |
|---|---|---|---|---|---|---|---|
| | $\mu_\infty$ (kPa) | $\alpha$ | $k_{11}$ (Pa.s) | $k_{21}$ (Pa.s) | $c_{21}$ | $\kappa$ (GPa) | $\rho$ (kg/m$^3$) |
| Deep gray matter | 2.33 | 4.92 | -1.68 | 647.65 | 0.62 | | |
| Cortical gray matter | 1.59 | -3.76 | -7.58 | 504.72 | 0.69 | | |
| Corpus Callosum | 2.07 | -2.32 | -0.29 | 453.03 | 0.59 | | |
| Corona Radiata | 2.08 | -3.47 | -7.72 | 754.340 | 0.66 | 2.19 | 1040 |
| Cerebellum gray matter | 1.68 | 2.71 | -0.45 | 379.20 | 0.61 | | |
| Cerebellum white matter | 2.13 | 2.71 | -1.43 | 586.09 | 0.62 | | |
| Brainstem | 2.27 | 2.61 | -0.58 | 578.85 | 0.61 | | |

**Table 6**. O-USS material properties of the various brain substructures for subject S03 (28/M).

| Brain substructure | Material properties (O-USS model) | | | | | | |
|---|---|---|---|---|---|---|---|
| | $\mu_\infty$ (kPa) | $\alpha$ | $k_{11}$ (Pa.s) | $k_{21}$ (Pa.s) | $c_{21}$ | $\kappa$ (GPa) | $\rho$ (kg/m$^3$) |
| Deep gray matter | 2.12 | 4.92 | -1.44 | 626.34 | 0.61 | | |
| Cortical gray matter | 1.62 | -3.76 | -3.41 | 508.12 | 0.66 | | |
| Corpus Callosum | 2.07 | -2.32 | -2.37 | 617.64 | 0.62 | | |
| Corona Radiata | 1.79 | -3.47 | -3.54 | 604.42 | 0.64 | 2.19 | 1040 |
| Cerebellum gray matter | 1.78 | 2.71 | 2.63e-04 | 397.85 | 0.59 | | |
| Cerebellum white matter | 1.81 | 2.71 | -0.69 | 479.05 | 0.60 | | |
| Brainstem | 1.68 | 2.61 | -0.32 | 436.19 | 0.60 | | |

### SM5. MPM simulation details

The Uintah software MPM package is used in this study to run computational simulations on the Blue Crab supercomputing cluster located at the Maryland Advanced Research Computing Center (MARCC). The segmented 3D voxelated brain image (1.5 mm isotropic resolution) from MRI and SWI is input as material points (see model illustration in Fig. 1(e)). A background grid size of 1.5 mm$^3$ is chosen, which translates to a 1 particle-per-cell configuration, in which each grid cell contains a single material point (or particle). The convected particle domain interpolation (CPDI) scheme[82] is employed to project particle information to the background grid. Explicit time integration is used along with the update stress last (USL) approach to advance the system to the next time step. Note that Uintah software implements the Fluid implicit particle (or FLIP) method for updating particle velocity and position (with first order approximation) from the grid. Further, a no-slip no-interpenetration contact is assigned between adjacent brain substructures or regions. The O-USS model (Eqs. (6-9)) was implemented in the Uintah software; model verification was conducted under multiple homogeneous deformation modes (viz., uniaxial tension and compression, simple and pure shear, equibiaxial tension, and hydrostatic compression) and at different strain rates (0.001-200 $s^{-1}$). Model parameters listed in Table 3 were used for the representative S1 subject, along with the rotational boundary condition in Fig. 1(f) (peak acceleration of approximately -227 rad/$s^2$, initial angular velocity of 3.42 rad/s). Simulated strain response is extracted at a 3 ms temporal resolution.

### References

1. Holzapfel, G.A. (2007). Nonlinear Solid Mechanics: A Continuum Approach for Engineering., 455.

2. Zhurov, A.I., Limbert, G., Aeschlimann, D.P., and Middleton, J. (2007). A constitutive model for the periodontal ligament as a compressible transversely isotropic visco-hyperelastic tissue. Comput. Methods Biomech. Biomed. Engin. 10, 223–235.
